# Controversy over the decline of arthropods: a matter of temporal baseline?

**DOI:** 10.1101/2022.02.09.479422

**Authors:** François Duchenne, Emmanuelle Porcher, Jean-Baptiste Mihoub, Grégoire Loïs, Colin Fontaine

## Abstract

Recently, a number of studies have reported somewhat contradictory patterns of temporal trends in arthropod abundance, from decline to increase. Arthropods often exhibit non-monotonous variation in abundance over time, making it important to account for temporal coverage in interpretation of abundance trends, which is often overlooked in statistical analysis. Combining four recently analysed datasets that led to contrasting outcomes, we first show that temporal abundance variations of arthropods are non-monotonous. Using simulations, we show non-monotony is likely to bias estimated linear abundance trends. Finally, analysing empirical data, we show that heterogeneity in estimated abundance trends is significantly related to the variation in temporal baseline of analysed time series. Once differences in baseline years, habitats and continents are accounted for, we do not find any statistical difference in estimated linear abundance trends among the four datasets. We also show that short time series produce more stochastic abundance trends than long series, making the dearth of old and long-term time series a strong limitation in the assessment of temporal trends in arthropod abundance. The lack of time series with a baseline year before global change acceleration is likely to lead to an underestimation of global change effects on biodiversity.

## Introduction

Over the last decades, many studies have documented a biodiversity crisis on the basis of high extinction rates (Dirzo & Raven 2003; Ceballos *et al*. 2015) and population losses or declines (Butchart *et al*. 2010; Ceballos *et al*. 2017), mainly in vertebrates. Worldwide erosion of biodiversity is caused by anthropogenic global change (Sala *et al*. 2000), i.e. a set of pressures including among others land use change, climate warming, overexploitation, or pollution (Steffen *et al*. 2006). In recent years, invertebrates, and especially arthropods, have been at the centre of a debate (Dornelas & Daskalova 2020; McDermott 2021) regarding the magnitude and even the directionality of the temporal trends in their abundance. Some studies showed a strong decline on the basis of standardized inventories (Hallmann *et al*. 2017; Seibold *et al*. 2019), while analyses of meta-datasets assembling heterogeneous time series evidenced a decline of terrestrial, but an increase of aquatic arthropods (Pilotto *et al*. 2020; van Klink *et al*. 2020) or no overall decline (Crossley *et al*. 2020). Finally, an analysis of opportunistic occurrence data revealed non-monotonous dynamics and no overall decline (Outhwaite *et al*. 2020).

Heterogeneity in population trends among studies is not surprising but the underlying causes need to be differentiated, particularly to tell apart abiotic/biotic factors from methodological factors. The dynamics of biodiversity changes are well-documented and their heterogeneity relatively well understood for vertebrates (Antão *et al*. 2020; Daskalova *et al*. 2020b, a; Leung *et al*. 2020) or for some specific functional groups of invertebrates, such as pollinators (Grab *et al*. 2019; Duchenne *et al*. 2020; Soroye *et al*. 2020; Millard *et al*. 2021). However, the heterogeneity in population trends remains poorly explained for most invertebrates. Several reasons may explain the contrasting patterns revealed by the studies involved in the arthropod decline debate. Global change pressures can vary in space among locations or ecological habitats, and species abilities to respond to environmental changes may depend on their traits or their evolutionary history (Helmus *et al*. 2010; Grab *et al*. 2019); hence spatial, ecological and taxonomic coverages are obvious sources of heterogeneity among studies and are widely discussed in the recent literature (Blowes *et al*. 2019; Pilotto *et al*. 2020). For example, some studies used global datasets with a bias towards the northern hemisphere (van Klink *et al*. 2020) while others considered national or even more local spatial extents (Hallmann *et al*. 2017). Some focused on terrestrial arthropods (Hallmann *et al*. 2017; Seibold *et al*. 2019) while others included aquatic groups (Crossley *et al*. 2020; van Klink *et al*. 2020). Finally, some studies analysed average trends by pooling multiple taxonomic groups (Hallmann *et al*. 2017; van Klink *et al*. 2020), while others reported trends for each taxonomic group (Outhwaite *et al*. 2020).

Differences in temporal coverage (*i*.*e*. the time period during **which a taxon or community was monitored**) across studies are at the core of the debate over biodiversity crisis (Pauly 1995; Cardinale *et al*. 2018; Loreau *et al*. 2022), but efforts to account for temporal coverage in statistical analysis remain limited (Cardinale *et al*. 2018; Didham *et al*. 2020; Loreau *et al*. 2022). Heterogeneity in temporal coverage is likely to influence the conclusions of studies assessing temporal trends in arthropod abundance for two reasons. First, there is variation in baseline years across the studies fuelling the debate: some analysed time series of identical temporal coverage and relatively old baseline year (Hallmann *et al*. 2017; Outhwaite *et al*. 2020), while others used heterogeneous datasets consisting mostly of time series with recent baseline years (Crossley *et al*. 2020; van Klink *et al*. 2020). Second, non-monotonous dynamics repeatedly reported in arthropods (Macgregor *et al*. 2019; Baranov *et al*. 2020; Høye *et al*. 2021; Schowalter *et al*. 2021), due to temporally variable environmental pressures (Baranov *et al*. 2020; Duchenne *et al*. 2020; Schowalter *et al*. 2021), can affect linear estimates of abundance trend beyond the original description of the shifting baseline syndrome (Fig. 1). With non-monotonous dynamics, estimated linear trends can vary from positive to negative when considering different baseline years for the same time series (e.g. Fig. 1h). As most arthropod trends were estimated assuming linear trends and available data often do not make it possible to account for all factors influencing temporal variation in abundance, it appears critical to account for differences in baseline years when making comparison within and among studies.

**Fig. 1:**
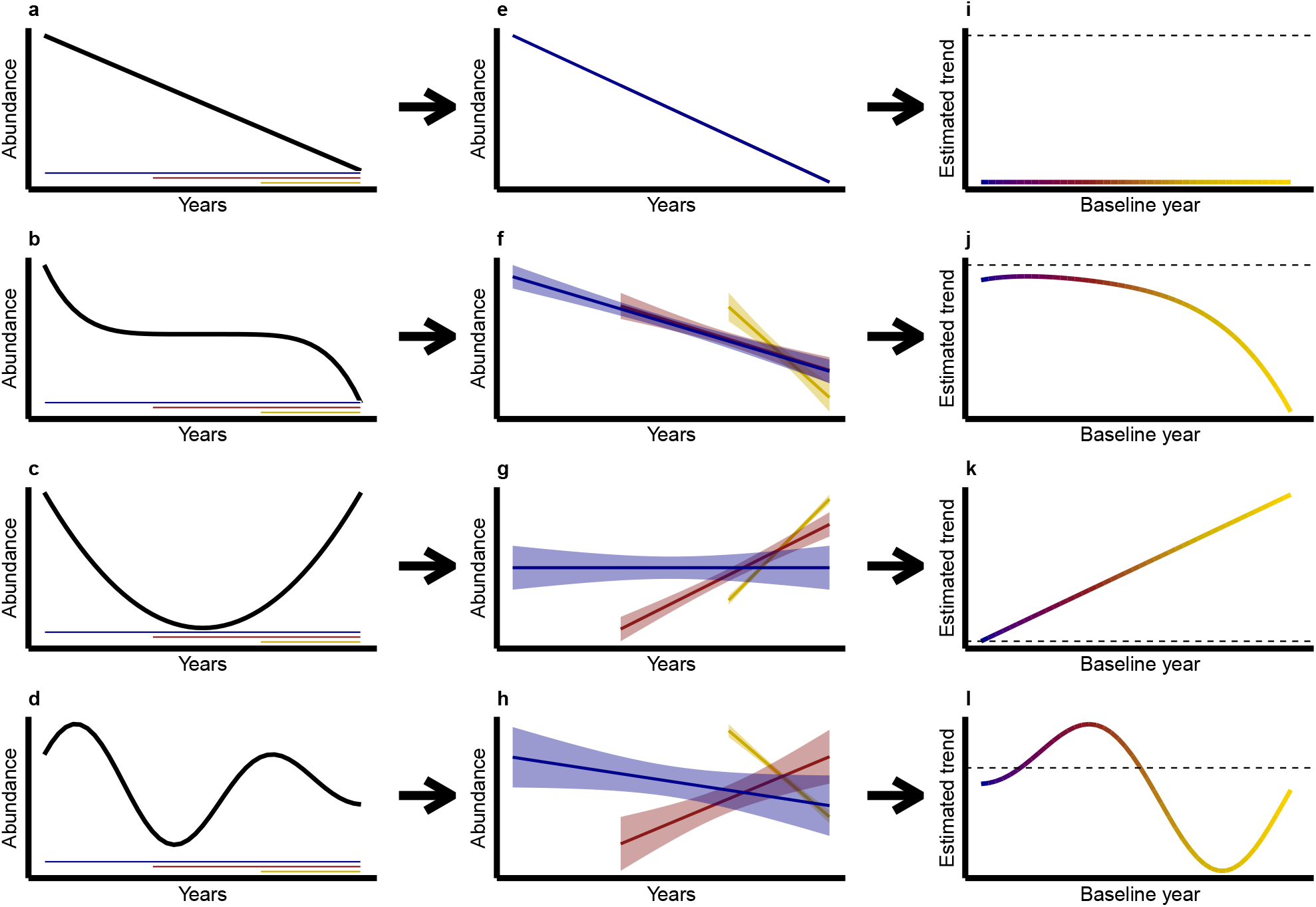
Schematic examples of how baseline year may affect estimated linear abundance trends. (a-d) Hypothetical time-series showing abundance variations over years (black) and three possible temporal coverages with contrasting baseline years: old (blue), intermediate (red) and recent (yellow). (e-h) Linear abundance trends estimated with time series starting from these three baseline years. (i-l) Pattern of estimated abundance trends against baseline year, reconstructed by estimating abundance trends over all possible baseline years. The dashed black line shows the zero-value delimiting positive (above) vs. negative (below) abundance trends.

Here, we evaluated the effect of the baseline years of time series on estimated linear trends in arthropod abundance, using the four largest datasets of arthropod time series recently analysed in studies fuelling the current debate (Dornelas *et al*. 2018; Crossley *et al*. 2020; Outhwaite *et al*. 2020; van Klink *et al*. 2020). We first characterized non-monotony in temporal variations of arthropod abundance. Then, using simulations we assessed how non-linearity can bias linear abundance trends. Finally, using a sliding baseline method on the empirical dataset aggregated here (*cf*. Methods, Fig. 2), we measured how non-linearity in arthropod abundance trends can produce a statistical dependency of estimated linear abundance trends on the baseline year.

**Fig. 2:**
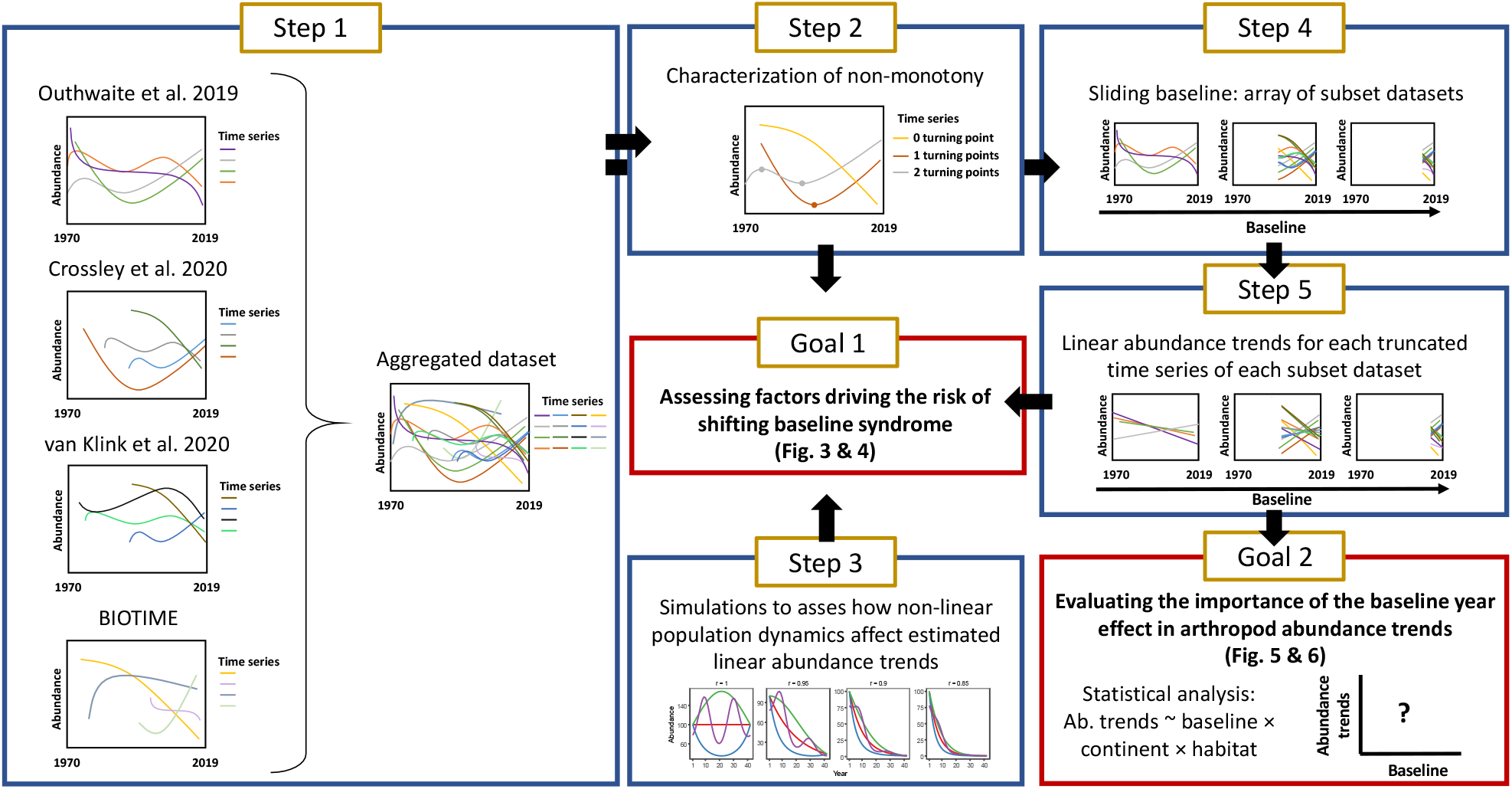
Schematic description of data aggregation, and workflow of analyses to assess the effect of the temporal baseline on abundance trends. In Step 1, we aggregated empirical datasets from four different source datasets. Then we characterized non-monotony of abundance variations over time (Step 2), and we assessed how non-monotony can affect estimated linear abundance trends using simulations (Step3, Goal 1). Finally, we created an array of subset datasets using a sliding baseline (Step 4) to estimate how baseline years influenced estimated linear abundance trends in empirical data (Step 5, Goal 2). Abundance trends are expressed as growth rates for all time series, allowing comparisons among common and rare species, as well as among datasets (cf. Supplementary Methods, Fig. S2 & S3). Statistical analyses were then performed accounting for habitat (aquatic vs terrestrial), continent, data source and taxonomy.

## Methods

### The four source datasets

We aggregated four source datasets from recent publications evaluating abundance trends in arthropods (Table S1): (i) annual occupancy estimates (the proportion of 1km^2^ grid cells in a region occupied by a species, a proxy for abundance) at species level for a wide diversity of arthropods from Great-Britain, produced by Outhwaite *et al*. (2019), (ii) annual estimates of arthropod abundances mostly at species level from American Long-Term Ecological Research (LTER) sites from Crossley *et al*. (2020), (iii) annual estimates of arthropod abundances from the meta-analysis of van Klink et al. (2020), aggregated at the resolution of taxonomic order, and (iv) abundance time series from the BIOTIME database (Dornelas *et al*. 2018), mostly at the species level (Fig. S1).

We focused on well covered continents and habitats, using only arthropod time series from North America and Europe with information on habitat (aquatic *vs*. terrestrial). We homogenized taxonomy using the Global Biodiversity Information Facility (GBIF) taxonomy backbone. For data from Crossley *et al*. (2020) and from van Klink *et al*. (2020), some time series describe the temporal variations of a wide diversity of species pooled together by summing their respective abundances. For these datasets, we removed all time series with taxonomic resolution coarser than taxonomic order, except for non-insect arthropods that are often grouped at taxonomic class level in available datasets (*Chilipoda, Diplododa, Collembola, Branchiopoda* etc.). We retained these groups and consider them to be the same rank as taxonomic orders in the following for simplicity. Details about aggregation and filtering step are available in Supplementary Methods.

To study baseline year effects on linear abundance trends estimated from time series with consistent ending dates, we removed time series ending before 2005 (n = 14,717). We further removed the few abundance values before 1970 (n = 1,039, 0.4%) to focus on the time period when most of the data were collected. Finally we removed time series shorter than 3 years (n = 47). This led to 14,130 original time series (Table S2, Fig. S1).

### Assessing the monotony of temporal variation in abundance (step 2, goal 1)

Because estimated linear abundance trends depend on the baseline year only if abundance varies non-linearly, especially non-monotonically, over time (Fig. 1), we estimated non-monotony using a Generalized Additive Model (GAM) for each original time series. We estimated the strength of non-monotony as the number of local extrema, hereafter turning points, observed in the non-linear trend predicted by the GAM, as detailed in supplementary Methods.

### Simulations of temporal declines for different population dynamics (step 3, goal 1)

We used simulations to assess how non-linearity in temporal variation of abundance (*i*.*e*. population dynamics) can bias estimated linear abundance trends, depending on their baseline year. We used four different shapes of population dynamics (Fig. 4a), using four functions describing temporal variation in average population size over 42 years (cf. R script available in supplementary materials). For each shape of population dynamics, we simulated time series with different growth rates *r* from stable (*r* = 1) to declining (*r* = [0.95, 0.9, 0.85]). The three latter values of *r* correspond to declines of 5%, 10% and 15% per year, respectively. We simulated 100 abundance time series for each shape of population dynamics and growth rate. To do so, for each year, we sampled observed abundance values from a Poisson distribution with a mean equal to the average population size of the corresponding year. Then, we estimated a linear abundance trend over the entire time period (baseline year at *t* = 1), as well as over truncated time series with different baseline years (*t* =10, 20, 30), using a Generalized linear model (GLM) with a Poisson error structure and a log link function, and accounting for temporal autocorrelation (equation (1) below).

### Generating an array of subset datasets using a sliding baseline (step 4)

To study the effect of the baseline year of time series on abundance trends estimated from the 14 130 empirical original time series, we created an array of 41 datasets, hereafter called subset datasets, corresponding to 41 different baseline years, from 1970 to 2010 by steps of one year (Fig. 2). For each of the subset datasets, time series were either truncated to start at the given baseline year, or removed if they did not include this specific baseline year. By construction, each of the original time series therefore appears several times in the array of 41 datasets, corresponding to *n* (1 ≤ *n* ≤ 41) truncated time series. Since time series with old baseline years are rare, the number of time series included in the subset datasets decreases with earlier baseline years.

### Estimating abundance trends (step 5)

We estimated arthropod abundance trends using one GLM per truncated time series. We considered only truncated time series with at least three annual estimates of abundance, including the abundance estimate in the year used to truncate the time series.

To obtain comparable abundance trends among the various sources, expressed in the same unit, we used a model structure that allows the estimation of growth rates. To do so we used GLMs with Poisson error structure with a *log* link function for count data, from van Klink *et al*. (2020), Crossley *et al*. (2020) and BIOTIME, GLMs with a binomial error structure with a *logit* link function for occupancy estimates from Outhwaite *et al*. (2019), and GLMs with a gaussian error structure with a *log* link function for density estimates from BIOTIME. Trends estimated with a *log* or *logit* link functions are expressed as the logarithm of a growth rate (see Supplementary Methods, Fig. S2 & S3), allowing comparisons between common and rare species, but also among datasets. Therefore, this approach with appropriate link functions gives the same importance to rare and to common species, in contrast to classic standardization of abundance by mean and standard deviations, which biases average trends by giving more weight to species with lower inter-annual variability in abundance (Fig. S4).

The GLM used for each truncated time series explains the abundance estimates of each year *i* by a Poisson or Binomial distribution, of parameter *λ*_*i*_ and *p*_*i*_ respectively, which depend on a linear year effect (*β*):

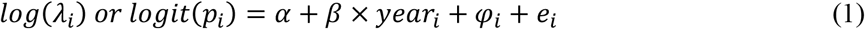

where *α* is the intercept, *φ*_*i*_ is a temporal random walk of order one 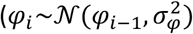, with *φ*_1_ = 0) to account for temporal autocorrelation and *e*_*i*_ an error term. We fitted these GLMs using the *INLA* R package (Rue *et al*. 2009).

### Evaluating the importance of the effect of baseline year in arthropod abundance trends (goal 2)

Pooling together the slopes of the year effect from the linear models presented above, across the 41 subset datasets, we obtained 192,244 abundance trends. We removed abundance trends estimated from time series with a single non-zero yearly estimate of abundance. We did so because growth rates estimated from such time series are likely to be extreme (*i*.*e*. strongly negative or positive) if this positive abundance estimate is by chance at the end or at the beginning of the truncated time series (Fig. S5). We thus kept the 175,796 abundance trends derived from truncated time series with ≥2 non-zero abundance estimate.

We examined the effect of baseline year used to truncate the original time series on abundance trends. To this end, we used a Bayesian linear mixed-effects model explaining abundance trends with spatial and temporal variables. Since we do not expect a linear relationship between abundance trends and baseline year (Fig. 1), the effect of baseline year was modelled as a polynomial of order three. We added a continent effect, a habitat effect, and all two-and three-way interactions between these effects and the polynomial baseline year effect (equation (2)), because recent results suggest that terrestrial and aquatic arthropods exhibit differences in their abundance trends and because population trends can vary over space (Outhwaite *et al*. 2020; van Klink *et al*. 2020). To assess whether differences among source datasets (Crossley *et al*. 2020; Outhwaite *et al*. 2020; van Klink *et al*. 2020) persist after spatio-temporal variables have been taken into account, we included a dataset effect, with four levels.

We also accounted for pseudo-replication in species belonging to the same taxonomic orders, as well as for the fact that several truncated time series are obtained from the same original time series, by adding a random effect of taxonomic order and time series ID on the intercept. Data from van Klink *et al*. (2020) and Crossley *et al*. (2020) originated from multiple sites and different data sources. Thus, we accounted for this structure by adding a random site effect nested in a random data source effect. For data from Outhwaite *et al*. (2019), the data source corresponds to the groups used to calculate occupancy estimates in the original dataset and the site corresponds to the country coverage of the data for each group (UK or GB), both extracted from the *Online-only Table 1* (Outhwaite *et al*. 2019).

The linear mixed-effect model is thus the following:

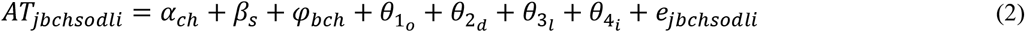

where *AT*_*jbchsodli*_is the abundance trend of truncated time series *j* with baseline *b*, from continent *c*, habitat *h*, dataset *s*, order *o*, data source *d*, site *l*, from original time series *i. α*_*ch*_ is the intercept for all combination of continent *c* and habitat *h*, while *β*_*s*_ is the effect of the source dataset *s. φ*_*bch*_ is baseline effect that depends on baseline *b*, continent *c* and habitat *h*. It is modelled as a temporal random walk of order one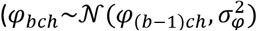, with *φ*_1*ch*_ = 0). The *θ*s denote random effects on the intercept: 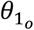 for taxonomic order, 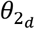 for data source, 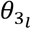 for site and 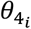 for time series ID. *e*_*jbchsodli*_is an error term, independent and identically distributed, following *𝒩*(0, *σ*^2^).

To account for the fact that our response variable is estimated, and thus each value has an associated standard error (*sde*_*j*_), we modelled the residual variance as a function of this error. In addition, since the four source datasets have different taxonomic scopes or different spatial scales, we expect that residual variance (*σ*^2^) will be strongly structured by the source dataset. Finally, the baseline year also affects the number of abundance estimates in time series, which is expected to affect the stochasticity of abundance trends (Bahlai *et al*. 2021). We modelled the dependence of variance of the residuals (*σ²*) on standard error associated to abundance trends of truncated time serie *j* (*AT*_*j*_), on source dataset *s* and on the number of years with data in each truncated time series (*ny*):

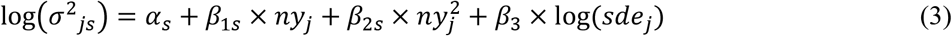

where *α*_*s*_ is the intercept, which depends on source dataset *s*, and *β*_1*s*_ and *β*_2*s*_ the polynomial effects of the number of years with data in truncated time series *j*, for each dataset, on the residual variance *σ*^2^_*sj*_. *β*_3_ is the effect of standard error associated to *AT*_*j*_ on the residual variance *σ*^2^_*sj*_. Parameters of the two models, one for the mean (equation (2)) and one for the residual variance (equation (3)), are estimated simultaneously. On this model (equation (2)), we estimated the variance of abundance trends explained by each random or fixed effect, as the ratio of its variance on the sum of all these variances plus the variance of the residuals. Priors used are detailed in the R script available in supplementary material. The model was fitted using the *R2jags* R package (Su & Yajima 2012), with 3 chains using 60,000 iterations with a burnin of 50,000 and a thin rate of 3, which was enough to reach convergence (all parameters with Rhat<1.1).

## Results

The number of turning points per empirical time series increases with the number of years with data in the time series, indicating that the vast majority of population trends are non-monotonous when time series are long enough (Fig. 3a). As expected from Figure 1, the strength of the non-monotony, measured as the number of turning points per time series, affects trend estimation. Non-monotonous abundance trends are characterized by multiple changes in trend direction over time, such that estimated linear trends may have opposite signs depending on the baseline year considered as the start of the time series (Fig. 3b). This pattern is consistent across the four source datasets (Fig. S6).

**Fig. 3:**
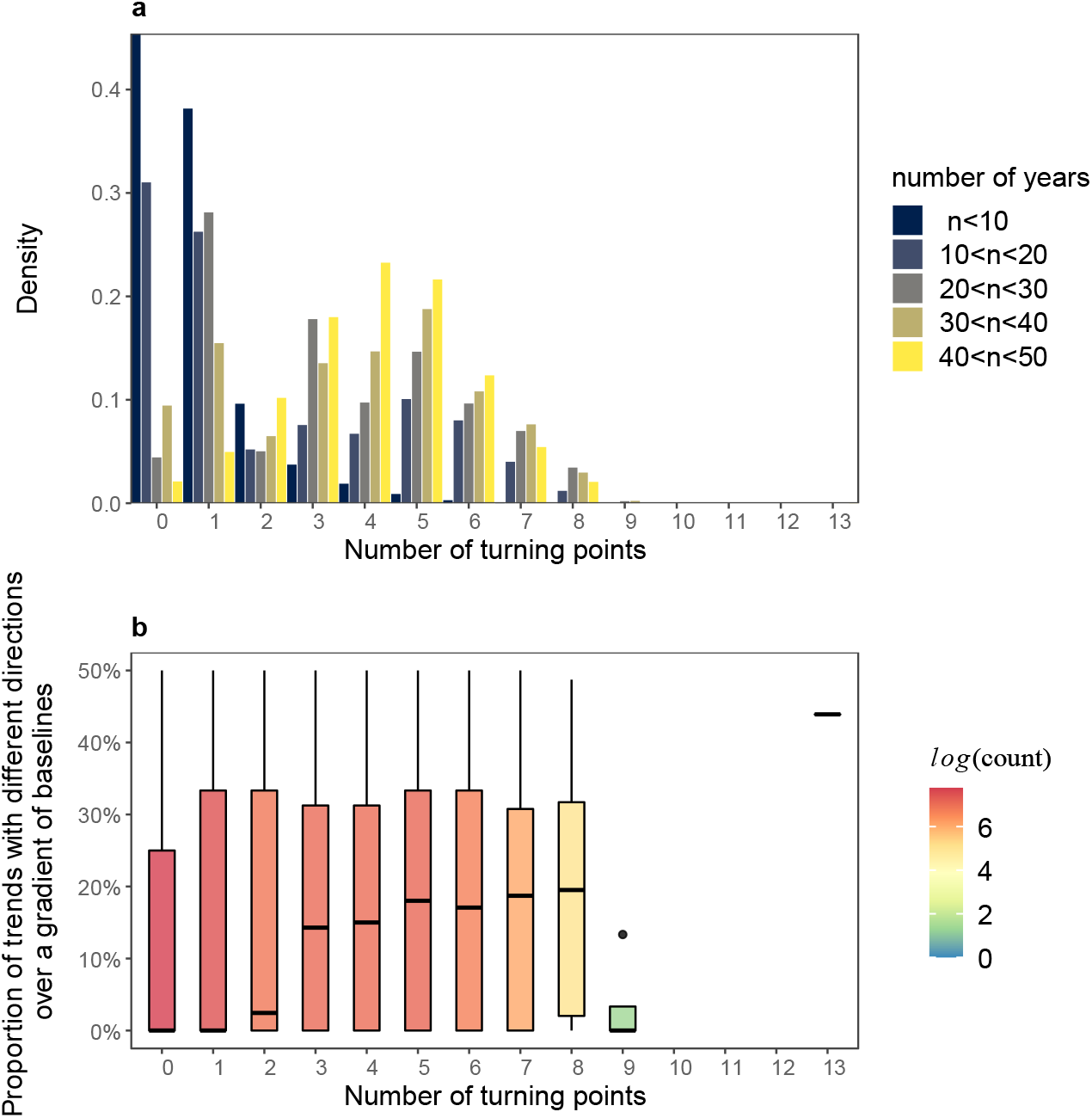
Empirical time series of arthropod abundance are non-monotonous, leading to unstable abundance trends over a gradient of baseline years. (a) Distribution of the number of original time series as a function of the number of turning points (a proxy for non-monotony, cf. Methods) and of the number of years with data in the corresponding time series. (b) Variation in direction of abundance trends (positive vs negative) of truncated time series, as a function of the number of years with data in the corresponding time series. Truncated time series are from the same original time series which is truncated for every possible baseline year (cf. Methods, Fig. 2). Boxplots represent minimum and maximum values (bottom and top of vertical lines), first and third quartiles (Q1 and Q3, bottom and top of boxes) and median (thick horizontal lines); colours indicate sample size (number of original time series). Points with values outside of the range [Q1-1.5(Q3-Q1), Q3+1.5(Q3-Q1)] are considered as outliers and represented as full circles.

Using simulated population dynamics (Fig. 4a), we show that similar rates of declines lead to different estimated growth rates, depending on the shape of the population dynamics (Fig. 4a-c). Assuming linearity to estimate population trends from non-linear dynamics can induce a strong bias when the growth rate departs from stability (*r*=1), by either overestimating or underestimating the genuine simulated trends (Fig. 4c). As expected, truncating the time series towards more recent baseline years increases the bias in the estimated growth rate, except in simulations with linear population dynamics (Fig. 4d). In addition to the magnitude of the bias, uncertainty (i.e. dispersion of values around the median) also increases when truncating the time series towards more recent baseline years (Fig. 4d). Short and recent time series tend to produce extreme estimated growth rates, which can even be opposite to the simulated long-term decline (Fig. 4d). These results suggest that not accounting for non-linearity in population trends can induce strong biases, and that short-term and recent time series cannot be used to infer long-term population change, even if population dynamics are linear, due to the strong uncertainties on estimated values.

**Fig. 4:**
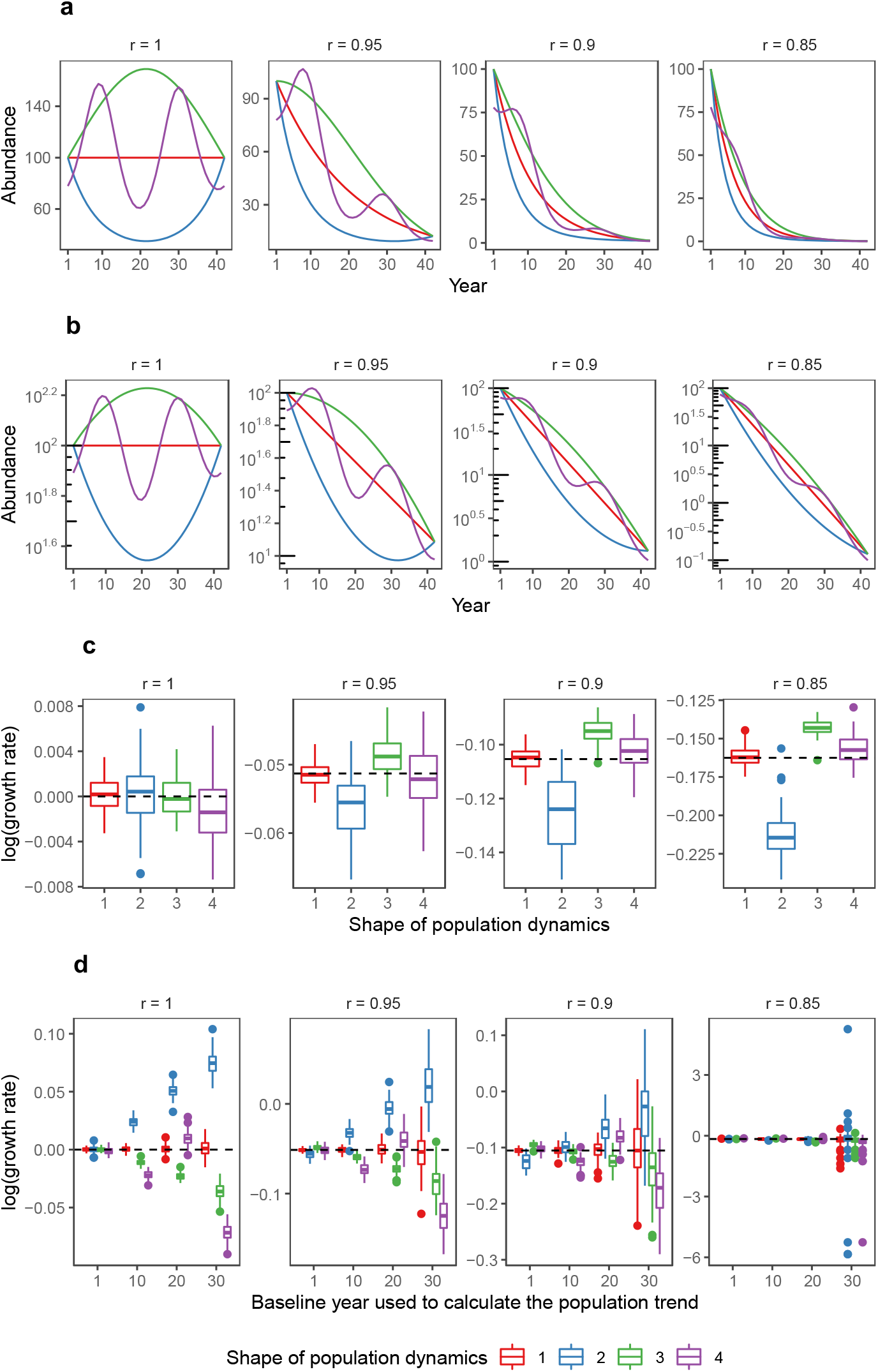
Non-linear population dynamics can bias estimated abundance trends. (a) Raw patterns of population dynamics when the population is stable, i.e. with a growth rate r = 1, or when the population declines by 5% (r=0.95), 10% (r=0.9) or 15% (r=0.85) each year. (b) Scaled patterns of population dynamics shown in (a) but using a log10 y-axis scale illustrating that the red curve corresponds to linear dynamics (cf. Supplementary methods, Fig. S2). (c) Estimated values of log(r) as a function of the shape of the population dynamics over 100 stochastic simulations per type of population dynamics. (d) Estimated values of log(r) as a function of the shape of the population dynamics and of the baseline year used to truncate the time series. In (c) and (d) the dashed horizontal line shows the value of the logarithm of the true (simulated) growth rate. Boxplots have the same meaning as in Figure 3. Right panel of (d) without outliers is represented in Figure S7.

When looking for general patterns in all empirical truncated time series together, statistical analyses show that average linear abundance trends estimated from the truncated time series strongly depend on the baseline year used for truncation, in interaction with habitats and continents (Fig. 5). This suggests that the non-linearity in temporal variations of abundance (Fig. 3b) leads to a strong dependence of the Arthropod population trends to the considered period (Fig. 5a). Results show a higher uncertainty in North America likely because time series with old baseline years are scarce for this region (Fig. 5b). This highlights that data are missing to estimate long-term abundance changes of Arthropods, and that recent data cannot help to fill this blank because of non-linearity.

**Fig. 5:**
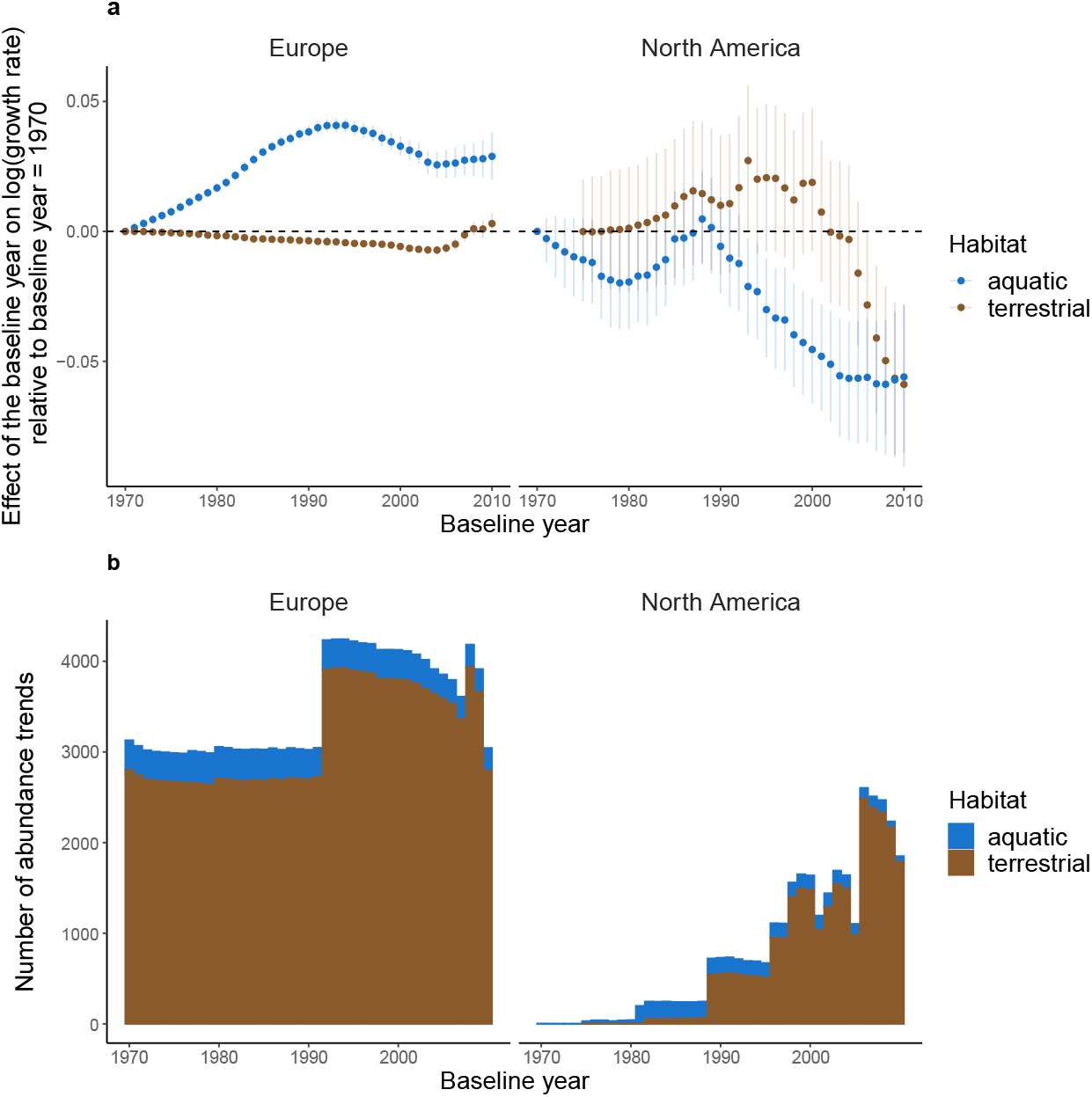
Effect of the baseline year on average abundance trends. (a) Relationships between the baseline year of times series and estimated abundance trends (log of the growth rate) relative to the abundance trends estimated with the oldest baseline of the dataset (1970), for each continent and habitat. Error bars are 95% confidence intervals. (b) Number of truncated time series used for each baseline value.

Importantly, for the two source datasets including short and long time series (*i*.*e*. those from Outhwaite *et al*. and van Klink *et al*., Fig. 6a), the residual variance of the model strongly decreases with the number of years with data in time series. This is consistent with our previous results based on simulations of truncated time series (Fig. 4c), and suggests that the stochasticity in abundance trends estimated from short time series is much greater than that from long time series. For data from Crossley *et al*. and BIOTIME, the relationship between the residual variance in abundance trends and the number of years with data is strongly parabolic. However, those datasets are strongly biased towards recent and short time series (Fig. 6a), prompting caution in interpreting this signal.

**Fig. 6:**
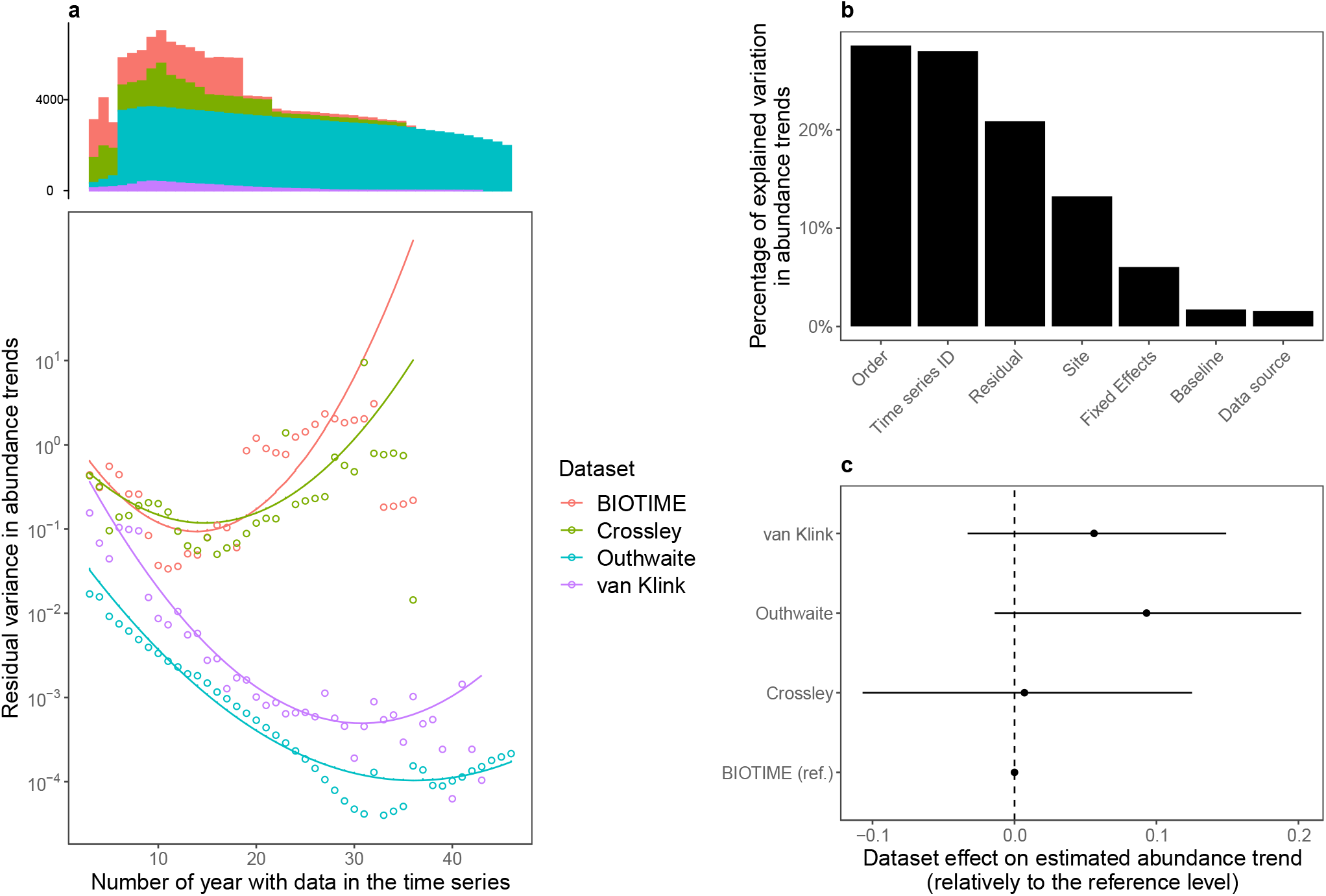
Variation in abundance trends is strongly structured by methodological effects. (a) Predicted (lines) and observed (points) residual variance of the model for each source dataset. Predictions are from equation (3). The histogram at the top shows the distribution of truncated time series along the x-axis, per dataset. (b) Variation in abundance trends explained by the model effects, and residual variation. (c) Black dots show the effect (± CI_95%_) of the source dataset on abundance trends, relatively to the reference level (BIOTIME).

Although statistically significant, the effects of baseline year explain only a small fraction of the total variation in abundance trends (Fig. 6b). Overall, random effects controlling for methodological issues, spatial and taxonomic heterogeneity explain 56% of the variation in abundance trends (Fig. 6b). A high proportion of this variation in abundance trends is explained by the random effect of time series ID, controlling for the artificial dependencies among abundance trends generated by the truncation procedure (*cf*. Methods). Most of the remaining variation in abundance trends is explained by taxonomic order and local site random effects, highlighting that abundance trends are strongly heterogeneous among clades and among sites. This questions the relevance of global estimates and stressing the need to carefully identify the drivers of such heterogeneity. Some groups, such as Trombidiformes (mites), Orthoptera, Collembola and Isopoda, exhibit more positive average abundance trends while other groups, such as Archaeognatha (jumping bristletails), Hymenoptera, Coleoptera, Dermaptera and Blattodea are associated with more negative trends (Fig. S8). Strikingly, once the various sources of heterogeneity are controlled for, we do not find any significant difference among the four source datasets from which the abundance estimates were extracted (Fig. 6c).

## Discussion

### Accounting for differences in baseline years contributes to settling the debate on arthropod decline

Our analysis shows that the non-monotony of empirical abundance time series induces a strong dependency of estimated abundance trends to the baseline year, as well as a strong uncertainty in abundance trends from short time series. Our findings therefore bring statistical support to the fact that most of the available data regarding arthropods, which are biased towards recent and short time series, are not appropriate to extrapolate long-term population trends, as suggested recently in discussions over vertebrates and arthropods population trends (Daskalova *et al*. 2021; Loreau *et al*. 2022; Mehrabi & Naidoo 2022).

We show that baseline year is statistically linked to abundance trends, and we highlight that average linear abundance trends, if they make any sense, should be interpreted in the light of the temporal window covered by the analyzed time series. By statistically assessing the effect of shifting baseline, we provide a general picture of how abundance trends change as function of baseline year, which may help to reposition the findings of past and future studies in a broader context, and hopefully to make between-studies comparison easier. Importantly, once we controlled average temporal trends of studies by the various sources of heterogeneity we did not find any statistical difference among source datasets, suggesting that the contradiction among recent results regarding arthropod decline comes from methodological issues related to temporal coverage as well as taxonomical and geographical bias.

The importance to define temporal baseline and use common spatial yardsticks when evaluating temporal change was previously emphasized in the wider context of biodiversity decline (Lotze & Worm 2009; Mihoub *et al*. 2017; Cardinale *et al*. 2018; Stouffer *et al*. 2021), but here we bring further statistical support to this caveat. Indeed, although the importance of comparing results with common baselines is well known, the baseline effect is rarely explicitly accounted for in quantitative analysis (but see Macgregor *et al*. (2019) for an example), in contrast to other sources of heterogeneity in abundance trends (*e*.*g*. space and taxonomy). Although van Klink et al. (2020) tested the effect of the starting year by truncating their time series (see Fig. 3 of van Klink *et al*. 2020), they did not formally test for a baseline effect: discarding data earlier than a given baseline threshold, from 1960 to 2005, did not result in constraining the baseline years to be equal across all time series. Since their dataset was mainly composed of time series with a baseline year after 1990, discarding data before a giving baseline threshold did not affect the overall distribution of baseline years much, except for recent thresholds (post-1990).

We also show that shorter time series exhibit much more stochasticity (*i*.*e*. higher residual variance) than long term series, which increases the uncertainty of results from these short-term time series that are commonly used in assessing arthropod abundance trends (Seibold *et al*. 2019; Crossley *et al*. 2020; van Klink *et al*. 2020). In other words, our results show that short-term series need to be much more replicated than long-term time series to reach the same level of confidence in the results. This is consistent with previous trends assessment regarding moths in Great-Britain (Macgregor *et al*. 2019) and more generally with the fact that arthropod abundance trends estimated from short time series are strongly sensitive to year to year variations (Daskalova *et al*. 2021). Here we show that uncertainty decreases exponentially with the length of time series, highlighting the importance to maintain existing biodiversity monitoring schemes. Obtaining long enough time series is critical for assessing reliable abundance trends, which echoes hot and recent debates about abundance trends in vertebrates (Leung *et al*. 2022; Loreau *et al*. 2022).

### Limitations and future challenges

In addition to the dramatic lack of data for some regions of the world, our study suggests that available arthropod abundances suffer from temporal limitations that should be carefully kept in mind when assessing their trends. First, the significant effects of baseline year and study area (continent and local site) suggest that the estimation of the arthropod abundance trends suffers from large uncertainties, mainly due to the lack of historical data. Here, 1970 is the oldest baseline we can tackle with the data at hand, but it cannot be considered as a reference before the rise in global change pressures (Mihoub *et al*. 2017), leading to a likely underestimation of global change effects on arthropod abundance. Since we cannot go back in time to sample biodiversity, this issue will remain difficult to solve. Museum collections and other sources of historical data could help fill this gap, although extracting reliable information from such data is still challenging (Isaac *et al*. 2014; Bartomeus *et al*. 2019; Duchenne *et al*. 2020; Outhwaite *et al*. 2020). Whether or not scientists can manage to obtain the necessary data and apply relevant methods to effectively turn back the clock, our analysis stresses the critical need to maintain long-term monitoring and secure appropriate archiving of related data (Millar & Searcy 2019).

Second, we used the same datasets as previous studies, so our analysis does not introduce new elements to assess the reliability of these abundance trends with respect to potential biases related to spatial and taxonomic coverage or data quality (Desquilbet *et al*. 2020). For example, some of the time series incl ded in van lin et al.’s and Crossle et al.’s datasets ere rod ced b ex eri ental st dies manipulating environmental conditions likely to influence abundance trends (Desquilbet *et al*. 2020, 2021), leading to criticisms about the use of these time series to assess temporal trends (Cardinale *et al*. 2018). Similarly, using only or mostly data from research stations, such as Long-Term Ecological Research sites, could bias estimated abundance trends upward as these locations are often partially protected from disturbances. Our results show that local site strongly explains heterogeneity in abundance trends, which is consistent with the strong influence of local changes in environmental conditions on arthropod abundance trends (Seibold *et al*. 2019). This has consequences for the interpretation of global trends obtained from a non-representative sample of sites. This potential bias stresses the need for standardized protocols to monitor arthropod abundance in numerous sites, representative of the areas covered by different habitats and land-uses, to handle the diversity of anthropic pressures on biodiversity, some remaining restricted to particular areas while others apply widely over space. Monitoring schemes based on citizen sciences are one way to tackle this challenge (van Swaay *et al*. 2008; Jeliazkov *et al*. 2016), since they can produce protocoled or semi-protocoled datasets over a large set of habitats and landscapes, over seasons and years, at national or even continental scales. Citizen science monitoring schemes are often recent, but would be of considerable help to ensure long-term monitoring of species abundances, should they be maintained in the future.

Moreover, by expressing abundance trends as growth rates, we gave the same importance to rare species as to common species, which could be debated. Decline of extremely rare species could be poorly informative and less likely to affect ecosystem functioning than decline of common species, while rare species can exhibit extreme abundance trends (Fig. S9), thus affecting average abundance trends in a non-negligible way. On the other hand, rare species can contribute greatly to some biodiversity metrics, such as species richness, phylogenetic diversity or functional diversity, making it relevant to weigh them similarly as common species for some purposes. Consistent with previous comments (O’Hara & ot e 2010; Desquilbet *et al*. 2021), we show that transforming abundance counts with *log*(*x*+1) before statistical analysis of the data, as done by Crossley *et al*. 2020 and van Klink *et al*. 2020, instead of using model structures adapted to the data (*i*.*e*. GLM instead of linear models), can introduce an asymmetrical bias, flattening the abundance trends of rare species (Fig. S2). In a similar way, standardizing data by dividing by standard deviation also strongly biases the relative values of abundance trends among species (Fig. S4), stressing the need to explicitly test the effect of any transformations performed on the data.

Finally, the large variation in abundance trends across sites, taxonomic groups, habitats and continents brings into question the relevance of producing global multitaxon linear trends. Global multitaxon trends are likely to be disconnected from the ecological causes and consequences of biodiversity changes. Losses in one place or one taxonomic group cannot be balanced by gains in another place or taxonomic group. Losses and gains can have contrasting ecological and evolutionary consequences, that need to be assessed at relevant ecological scales, *e*.*g*. at community level or for a given functional group. Moreover, our results show that variation in arthropod abundance over time is non-linear and sometimes non-monotonous. This suggests that the use of linear analyses is inadequate, despite being the most straightforward and used analysis, and should at least always be associated to the temporal coverage of the data. Disconnecting arthropod decline assessments from temporal yardsticks can affect the understanding of published results making them apparently contradictory. This is particularly important for topics of interest for the general public such as arthropod decline, as it could lead to undermining trust in science (Dornelas & Daskalova 2020).

## Supporting information

Supplementary informations

## Acknowledgements

We thank all authors of previous studies on the topic, especially van Klink *et al*. (2020), Crossley *et al*. (2020), Outhwaite *et al*. (2019), Daskalova *et al*. (2021) and Dornelas *et al*. (2018) for making the data underlying their work available, and for building step by step our knowledge about the way biodiversity change over time. We also thank an anonymous reviewer from a previous submission for his/her insightful comments and opinions on that manuscript. The simulations were performed at the HPCaVe centre at Sorbonne Université. François Duchenne was funded by the European Research Council (ERC) under the E ro ean Union’s Hori on 0 0 researc and innovation rogra (grant agree ent N° 7876 8). Version 3 of this article (https://doi.org/10.1101/2022.02.09.479422) has been peer-reviewed and recommended by Peer Community In Ecology (https://doi.org/10.24072/pci.ecology.100098).

## Data, scripts and codes availability

All data used here were publicly available (cf. Methods). Scripts used for analyses are available in Supplementary materials (https://doi.org/10.1101/2022.02.09.479422) and here: https://github.com/f-duchenne/Controversy-over-the-decline-of-arthropods.

## Supplementary material

Supplementary material are available online: https://doi.org/10.1101/2022.02.09.479422

## References

Antão, L.H., Bates, A.E., Blowes, S.A., Waldock, C., Supp, S.R., Magurran, A.E., et al. (2020). Temperature-related biodiversity change across temperate marine and terrestrial systems. Nat. Ecol. Evol., 4, 927–933. https://doi.org/10.1038/s41559-020-1185-7

Bahlai, C.A., White, E.R., Perrone, J.D., Cusser, S. & Stack Whitney, K. (2021). The broken window: An algorithm for quantifying and characterizing misleading trajectories in ecological processes. Ecol. Inform., 64, 101336. https://doi.org/10.1016/j.ecoinf.2021.101336

Baranov, V., Jourdan, J., Pilotto, F., Wagner, R. & Haase, P. (2020). Complex and nonlinear climate-driven changes in freshwater insect communities over 42 years. Conserv. Biol., 34, 1241–1251. https://doi.org/10.1111/cobi.13477

Bartomeus, I., Stavert, J.R., Ward, D. & Aguado, O. (2019). Historical collections as a tool for assessing the global pollination crisis. Philos. Trans. R. Soc. B Biol. Sci., 374, 20170389. https://doi.org/10.1098/rstb.2017.0389

Blowes, S.A., Supp, S.R., Antão, L.H., Bates, A., Bruelheide, H., Chase, J.M., et al. (2019). The geography of biodiversity change in marine and terrestrial assemblages. Science, 366, 339–345. https://doi.org/10.1126/science.aaw1620

Butchart, S.H.M., Walpole, M., Collen, B., Strien, A. van, Scharlemann, J.P.W., Almond, R.E.A., et al. (2010). Global Biodiversity: Indicators of Recent Declines. Science, 328, 1164–1168. https://doi.org/10.1126/science.1187512

Cardinale, B.J., Gonzalez, A., Allington, G.R.H. & Loreau, M. (2018). Is local biodiversity declining or not? A summary of the debate over analysis of species richness time trends. Biol. Conserv., 219, 175–183. https://doi.org/10.1016/j.biocon.2017.12.021

Ceballos, G., Ehrlich, P.R., Barnosky, A.D., García, A., Pringle, R.M. & Palmer, T.M. (2015). Accelerated modern human-induced species losses: Entering the sixth mass extinction. Sci. Adv., 1, e1400253. https://doi.org/10.1126/sciadv.1400253

Ceballos, G., Ehrlich, P.R. & Dirzo, R. (2017). Biological annihilation via the ongoing sixth mass extinction signaled by vertebrate population losses and declines. Proc. Natl. Acad. Sci., 114, E6089–E6096. https://doi.org/10.1073/pnas.1704949114

Crossley, M.S., Meier, A.R., Baldwin, E.M., Berry, L.L., Crenshaw, L.C., Hartman, G.L., et al. (2020). No net insect abundance and diversity declines across US Long Term Ecological Research sites. Nat. Ecol. Evol., 4, 1368–1376. https://doi.org/10.1038/s41559-020-1269-4

Daskalova, G.N., Myers-Smith, I.H., Bjorkman, A.D., Blowes, S.A., Supp, S.R., Magurran, A.E., et al. (2020a). Landscape-scale forest loss as a catalyst of population and biodiversity change. Science, 368, 1341–1347. https://doi.org/10.1126/science.aba1289

Daskalova, G.N., Myers-Smith, I.H. & Godlee, J.L. (020b). Rare and common vertebrates span a wide spectrum of population trends. Nat. Commun., 11, 4394. https://doi.org/10.1038/s41467-020-17779-0

Daskalova, G.N., Phillimore, A.B. & Myers-Smith, I.H. (2021). Accounting for year effects and sampling error in temporal analyses of invertebrate population and biodiversity change: a comment on Seibold et al. 2019. Insect Conserv. Divers., 14, 149–154. https://doi.org/10.1111/icad.12468

Desquilbet, M., Cornillon, P.-A., Gaume, L. & Bonmatin, J.-M. (2021). Adequate statistical modelling and data selection are essential when analysing abundance and diversity trends. Nat. Ecol. Evol., 5, 592–594. https://doi.org/10.1038/s41559-021-01427-x

Desquilbet, M., Gaume, L., Grippa, M., Céréghino, R., Humbert, J.-F., Bonmatin, J.-M., et al. (2020). Comment on “Meta-analysis reveals declines in terrestrial but increases in freshwater insect abundances.” Science, 370. https://doi.org/10.1126/science.abd8947

Didham, R.K., Basset, Y., Collins, C.M., Leather, S.R., Littlewood, N.A., Menz, M.H.M., et al. (2020). Interpreting insect declines: seven challenges and a way forward. Insect Conserv. Divers., 13, 103–114. https://doi.org/10.1111/icad.12408

Dirzo, R. & Raven, P.H. (2003). Global State of Biodiversity and Loss. Annu. Rev. Environ. Resour., 28, 137–167. https://doi.org/10.1146/annurev.energy.28.050302.105532

Dornelas, M., Antão, L.H., Moyes, F., Bates, A.E., Magurran, A.E., Adam, D., et al. (2018). BioTIME: A database of biodiversity time series for the Anthropocene. Glob. Ecol. Biogeogr., 27, 760–786. https://doi.org/10.1111/geb.12729

Dornelas, M. & Daskalova, G.N. (2020). Nuanced changes in insect abundance. Science, 368, 368–369. https://doi.org/10.1126/science.abb6861

Duchenne, F., Thébault, E., Michez, D., Gérard, M., Devaux, C., Rasmont, P., et al. (2020). Long-term effects of global change on occupancy and flight period of wild bees in Belgium. Glob. Change Biol., 26, 6753–6766. https://doi.org/10.1111/gcb.15379

Grab, H., Branstetter, M.G., Amon, N., Urban-Mead, K.R., Park, M.G., Gibbs, J., et al. (2019). Agriculturally dominated landscapes reduce bee phylogenetic diversity and pollination services. Science, 363, 282–284. https://doi.org/10.1126/science.aat6016

Hallmann, C.A., Sorg, M., Jongejans, E., Siepel, H., Hofland, N., Schwan, H., et al. (2017). More than 75 percent decline over 27 years in total flying insect biomass in protected areas. PLOS ONE, 12, e0185809. https://doi.org/10.1371/journal.pone.0185809

Helmus, M.R., Keller, W. (Bill), Paterson, M.J., Yan, N.D., Cannon, C.H. & Rusak, J.A. (2010). Communities contain closely related species during ecosystem disturbance. Ecol. Lett., 13, 162–174. https://doi.org/10.1111/j.1461-0248.2009.01411.x

Høye, T.T., Loboda, S., Koltz, A.M., Gillespie, M.A.K., Bowden, J.J. & Schmidt, N.M. (2021). Nonlinear trends in abundance and diversity and complex responses to climate change in Arctic arthropods. Proc. Natl. Acad. Sci., 118. https://doi.org/10.1073/pnas.2002557117

Isaac, N.J.B., Strien, A.J. van, August, T.A., Zeeuw, M.P. de & Roy, D.B. (2014). Statistics for citizen science: extracting signals of change from noisy ecological data. Methods Ecol. Evol., 5, 1052–1060. https://doi.org/10.1111/2041-210X.12254

Jeliazkov, A., Bas, Y., Kerbiriou, C., Julien, J.-F., Penone, C. & Le Viol, I. (2016). Large-scale semi-automated acoustic monitoring allows to detect temporal decline of bush-crickets. Glob. Ecol. Conserv., 6, 208–218. https://doi.org/10.1016/j.gecco.2016.02.008

van Klink, R., Bowler, D.E., Gongalsky, K.B., Swengel, A.B., Gentile, A. & Chase, J.M. (2020). Meta-analysis reveals declines in terrestrial but increases in freshwater insect abundances. Science, 368, 417–420. https://doi.org/10.1126/science.aax9931

Leung, B., Hargreaves, A.L., Greenberg, D.A., McGill, B., Dornelas, M. & Freeman, R. (2020). Clustered versus catastrophic global vertebrate declines. Nature, 588, 267–271. https://doi.org/10.1038/s41586-020-2920-6

Leung, B., Hargreaves, A.L., Greenberg, D.A., McGill, B., Dornelas, M. & Freeman, R. (2022). Reply to: Do not downplay biodiversity loss. Nature, 601, E29–E31. https://doi.org/10.1038/s41586-021-04180-0

Loreau, M., Cardinale, B.J., Isbell, F., Newbold, T., O’Connor, M.I. & de Mazancourt, C. (2022). Do not downplay biodiversity loss. Nature, 601, E27–E28. https://doi.org/10.1038/s41586-021-04179-7

Lotze, H.K. & Worm, B. (2009). Historical baselines for large marine animals. Trends Ecol. Evol., 24, 254–262. https://doi.org/10.1016/j.tree.2008.12.004

Macgregor, C.J., Williams, J.H., Bell, J.R. & Thomas, C.D. (2019). Moth biomass has fluctuated over 50 years in Britain but lacks a clear trend. Nat. Ecol. Evol., 3, 1645–1649. https://doi.org/10.1038/s41559-019-1028-6

McDermott, A. (2021). News Feature: To understand the plight of insects, entomologists look to the past. Proc. Natl. Acad. Sci., 118. https://doi.org/10.1073/pnas.2018499117

Mehrabi, Z. & Naidoo, R. (2022). Shifting baselines and biodiversity success stories. Nature, 601, E17–E18. https://doi.org/10.1038/s41586-021-03750-6

Mihoub, J.-B., Henle, K., Titeux, N., Brotons, L., Brummitt, N.A. & Schmeller, D.S. (2017). Setting temporal baselines for biodiversity: the limits of available monitoring data for capturing the full impact of anthropogenic pressures. Sci. Rep., 7. https://doi.org/10.1038/srep46781

Millar, E. & Searcy, C. (2019). The presence of citizen science in sustainability reporting. Sustain. Account. Manag. Policy J.

Millard, J., Outhwaite, C.L., Kinnersley, R., Freeman, R., Gregory, R.D., Adedoja, O., et al. (2021). Global effects of land-use intensity on local pollinator biodiversity. Nat. Commun., 12, 2902. https://doi.org/10.1038/s41467-021-23228-3

O’Hara, R.B. & Kotze, D.J. (2010). Do not log-transform count data. Methods Ecol. Evol., 1, 118–122. https://doi.org/10.1111/j.2041-210X.2010.00021.x

Outhwaite, C.L., Gregory, R.D., Chandler, R.E., Collen, B. & Isaac, N.J.B. (2020). Complex long-term biodiversity change among invertebrates, bryophytes and lichens. Nat. Ecol. Evol., 4, 384–392. https://doi.org/10.1038/s41559-020-1111-z

Outhwaite, C.L., Powney, G.D., August, T.A., Chandler, R.E., Rorke, S., Pescott, O.L., et al. (2019). Annual estimates of occupancy for bryophytes, lichens and invertebrates in the UK, 1970-2015. Sci. Data, 6, 259. https://doi.org/10.1038/s41597-019-0269-1

Pauly, D. (1995). Anecdotes and the shifting baseline syndrome of fisheries. Trends Ecol. Evol., 10, 430. https://doi.org/10.1016/S0169-5347(00)89171-5

Pilotto, F., Kühn, I., Adrian, R., Alber, R., Alignier, A., Andrews, C., et al. (2020). Meta-analysis of multidecadal biodiversity trends in Europe. Nat. Commun., 11, 3486. https://doi.org/10.1038/s41467-020-17171-y

Rue, H., Martino, S. & Chopin, N. (2009). Approximate Bayesian inference for latent Gaussian models by using integrated nested Laplace approximations. J. R. Stat. Soc. Ser. B Stat. Methodol., 71, 319–392. https://doi.org/10.1111/j.1467-9868.2008.00700.x

Sala, O.E., Chapin, F.S., Iii Armesto, J.J., Berlow, E., Bloomfield, J., et al. (2000). Global Biodiversity Scenarios for the Year 2100. Science, 287, 1770–1774. https://doi.org/10.1126/science.287.5459.1770

Schowalter, T.D., Pandey, M., Presley, S.J., Willig, M.R. & Zimmerman, J.K. (2021). Arthropods are not declining but are responsive to disturbance in the Luquillo Experimental Forest, Puerto Rico. Proc. Natl. Acad. Sci., 118. https://doi.org/10.1073/pnas.2002556117

Seibold, S., Gossner, M.M., Simons, N.K., Blüthgen, N., Müller, J, Ambarli, D., et al (2019) Atrt ro od decline in grasslands and forests is associated with landscape-level drivers. Nature, 574, 671–674. https://doi.org/10.1038/s41586-019-1684-3

Soroye, P., Newbold, T. & Kerr, J. (2020). Climate change contributes to widespread declines among bumble bees across continents. Science, 367, 685–688. https://doi.org/10.1126/science.aax8591

Steffen, W., Sanderson, R.A., Tyson, P.D., Jäger, J., Matson, P.A., Iii, B.M., et al. (2006). Global Change and the Earth System: A Planet Under Pressure. Springer Science & Business Media. https://doi.org/10.1007/b137870

Stouffer, P.C., Jirinec, V., Rutt, C.L., Bierregaard, R.O., Hernánde Palma, A., Johnson, E.l., et al (2021) Long-term change in the avifauna of undisturbed Amazonian rainforest: ground-foraging birds disappear and the baseline shifts. Ecol. Lett., 24, 186–195. https://doi.org/10.1111/ele.13628

Su, Y. & Yajima, M. (2012). Package ‘R2jags’. A Package for Running jags from R.

van Swaay, C.A.M., Nowicki, P., Settele, J. & van Strien, A.J. (2008). Butterfly monitoring in Europe: methods, applications and perspectives. Biodivers. Conserv., 17, 3455–3469. https://doi.org/10.1007/s10531-008-9491-4

