## Supplementary informations for "Controversy over the decline of arthropods: a matter of temporal baseline?"

**This PDF file includes:**

Supplementary Methods

Fig. S1 to 9

Table S1 & S2

### Supplementary Methods

*Aggregating datasets*

For data from Outhwaite *et al.* (2019), we focused on occupancy estimates labelled at Great-Britain level, which can in fact correspond to two different spatial coverages (United Kingdom or Great-Britain). Since for this dataset occupancy estimates are modelled, not observed, we excluded occupancy estimates that converged poorly in their analyses (Rhat<1.2): this yielded 138,095 occupancy estimates for 3,496 species, from 1970 to 2015, which correspond to 57% of the arthropod occupancy estimates at Great-Britain level from the original dataset.

For data from Crossley *et al.* (2020) (https://datadryad.org/stash/dataset/doi:10.5061/dryad.cc2fqz645) we used the file "External_Database_S1_PerSpecies_Abundance_LTER_annotated.csv." After data filtering, we obtained 78,187 annual abundance estimates from 1943 to 2019 with a taxonomic resolution varying from *species* to *order* levels. This corresponds to 94% of the abundance estimates from the original dataset.

For data from van Klink *et al.* (2020) we used the file "aax9931-Van-Klink-SM-CORRECTED-Data-S1.txt." Some data from American LTER sites and from BIOTIME could be duplicated with time series from Crossley *et al.* (2020) and from BIOTIME respectively, but because van Klink *et al.* aggregated data at taxonomic order (Fig. S1), we considered them as different. We also removed biomass data to keep abundance data only. We restricted our analysis to time series from North America and Europe as other locations were poorly represented. After data filtering, we obtained 26,785 annual abundance estimates from 1932 to 2018 with a taxonomic resolution at *order* level. This corresponds to 43% of the abundance estimates from the original dataset.

For BIOTIME data we extracted all data, and we kept only time series regarding arthropods from terrestrial and freshwater habitats, from North America and Europe. We removed LTER data to avoid duplicated time series with Crossley *et al.* (2020). We inferred zeros on a given plot for a given species for a given year when the plot has data for at data from the same taxonomic order than the focal species for the given year. Then we removed time series with only zero abundance values. After data filtering, we obtained 79,250 annual abundance estimates from 1898 to 2016 with a taxonomic resolution varying from *species* to *order* levels.

*Assessing monotony of abundance variation over time*

We estimated non-monotony of temporal abundance variations using a Generalized Additive Model (GAM) for each original time series, using the *mgcv* R package (Wood 2017). These GAMs have a Poisson error structure (with a *log* link function) for abundance data from van Klink *et al.* (2020), Crossley *et al.* (2020) and BIOTIME, or a gaussian error structure for logit transformed occupancy estimates from Outhwaite *et al.* (2019). We used a smooth (spline) effect of the year. For time series sampled at different periods, we added a spline effect of the period penalized by a ridge penalty, to model it as a random effect. To avoid non-identifiable models, the dimension of the basis used to represent the year smooth term is constrained to be smaller than to the number of years (duration) of the time series, minus one if it is sampled over different sampling periods, with a maximum basis value of 10.

Then extracting the polynomial effect of the year on abundance, we assessed the non-monotony of this polynomial effect as the number of local maximums and minimums (*i.e.* turning points) observed.

*Comparing abundance trends among species*

A common problem when comparing abundance trends over many species is that estimated trends are not easily comparable among species, especially between rare and common species, since abundance trends depend on initial abundance. Indeed, two species, a rare and an abundant one, with an abundance shifting in a year from 10 to 15 individuals and from 10,000 to 15,000 individuals respectively, are both growing with a rate of 50% per year. However, measuring these abundance variations on the number of individuals will give a gain of 5 individuals for the first one against a gain of 5,000 individuals for the second species. Demographic effects measured on rough measure of abundance (number of individuals, occupancy probability, etc.) depend on the level of abundance, hiding the decline of rare species (Fig. S2 & S3). Note that standardizing abundances of each species by subtracting the mean and dividing by standard deviation does not solve this problem in a proper way, since it does not allow to estimate growth rate but can instead create artefacts by increasing the importance of stable species with low inter-annual variation in their abundance, relatively to species declining/increasing but with high inter-annual variations in abundance (Fig. S4).

This dependency of abundance trends to initial abundance can be overcome by expressing population trends in terms of growth rates, a multiplicative factor which correspond to the growth of the population. In a time-discrete system, a growth rate > 1 is associated with increasing abundance over time, a growth rate equals to 1 corresponds to a constant abundance over time, while a growth rate < 1 is associated with decreasing abundance over time. Such measure allows to compare trends among species, regardless their initial abundance, but is cannot be estimated directly using classic linear models. Indeed, the demography of a population with growth rate *r*, as shown in Fig. S6, can be expressed by the following geometric progression:

$$A_t = A_{t=0} \times r^t$$

where $A_t$ is the abundance at time *t*, $A_{t=0}$ is the initial abundance (intercept), while *r* is the growth rate. The problem is that this product cannot be estimated using classic linear models which, by definition, estimate linear functions ($y = b + ax$). Here, we transformed occupancy estimates using a *logit* function and abundance counts using a *log* function to linearize geometric changes, then allowing to estimate growth rates using linear models. If we study the logarithm of the abundance instead of the abundance, then the function describing the population demography becomes linear:

$$\log(A_t) = \log(A_{t=0} \times r^t)$$
$$\log(A_t) = \log(A_{t=0}) + \log(r^t)$$
$$\log(A_t) = \log(A_{t=0}) + t \times \log(r)$$

If we set $a = \log(r)$ and $b = \log(A_{t=0})$, we have:

$$\log(A_t) = b + at$$

Thus, by regressing linearly *log(abundance)* against time we can estimate the logarithm of the growth rate, which is a measure of abundance trend independent of the initial abundance (*i.e.* the rarity) of species (Figure S6). Since log(1) = 0, then the sign and the value of the estimated slope indicate the direction and the magnitude of the abundance trend, respectively. However, since *log*(0) is not defined and since we have zero abundance values, we need to use a model structure which allows zero abundances. Usually, studies use $\log(A_t + \varepsilon)$ instead of $\log(A_t)$, choosing $\varepsilon = 1$ arbitrarily (Crossley *et al.* 2020; van Klink *et al.* 2020), while the value of $\varepsilon$ will strongly affect the result by breaking the linearization of the geometric progression (Fig. S2).

Here, to avoid this arbitrary transformation, we used a Generalized linear model (GLM) with a Poisson error structure, using a $log$ link function, which allows to estimate $\log(A_t) = b + at$ by considering zero abundance data.

Regarding the occupancy probabilities, the same logic applies but we have to account for the fact that the proportion of grid cells occupied by the species is bounded between 0 and 1. So the population dynamic can be view as a logistic regression with a carrying capacity (*K*) of 1:

$$\frac{dP}{dt} = r^*P\left(1 - \frac{P}{K}\right)$$

Where $r^*$ is the growth rate of a time-continuous system (0 = stable, negative = decline, positive = increase). Thus, we have:

$$P_t = \frac{KP_0e^{r^*t}}{K + P_0(e^{r^*t} - 1)} = \frac{K}{1 + \frac{K - P_0}{P_0}e^{-r^*t}}$$

If we set *K* = 1, we have:

$$P_t = \frac{1}{1 + \frac{1 - P_0}{P_0}e^{-r^*t}}$$

If we logit transform $P_t$, we get:

$$\log\left(\frac{P_t}{1 - P_t}\right) = \log(P_t) - \log(1 - P_t) = -\log\left(\frac{K - P_0}{P_0}\right) + r^*t$$

Switching from $r^*$ to $r$, so from continuous to discrete time we get:

$$\log\left(\frac{P_t}{1 - P_t}\right) = -\log\left(\frac{K - P_0}{P_0}\right) + t \times \log(r)$$

Where $r$ is the growth rate of a discrete time system (1 = stable, below 1 = decline, above 1 = increase). If we set $a = \log(r)$ and $b = -\log\left(\frac{K-P_0}{P_0}\right)$, we have:

$$\log\left(\frac{P_t}{1 - P_t}\right) = b + at$$

Thus, by regressing linearly *logit(occupancy probability)* against time we can estimate, as previously, the logarithm of the growth rate, which is a measure of abundance trend independent of the initial abundance (*i.e.* the rarity) of species (Figure S3).

So, here we used a GLM with a binomial error structure, using a *logit* link function, which allows to estimate $\mathrm{logit}(P_t) = b + at$.

**References**

Crossley, M.S., Meier, A.R., Baldwin, E.M., Berry, L.L., Crenshaw, L.C., Hartman, G.L., *et al.* (2020). No net insect abundance and diversity declines across US Long Term Ecological Research sites. *Nat. Ecol. Evol.*, 4, 1368–1376.

van Klink, R., Bowler, D.E., Gongalsky, K.B., Swengel, A.B., Gentile, A. & Chase, J.M. (2020). Meta-analysis reveals declines in terrestrial but increases in freshwater insect abundances. *Science*, 368, 417–420.

Wood, S.N. (2017). *Generalized Additive Models: An Introduction with R*. 2nd edn. Chapman and Hall/CRC, Boca Raton.

***Fig. S1: Taxonomic distribution of time series.*** *(a) Number of species across taxonomic groups and (b) source datasets, as a function of their taxonomic resolution.*

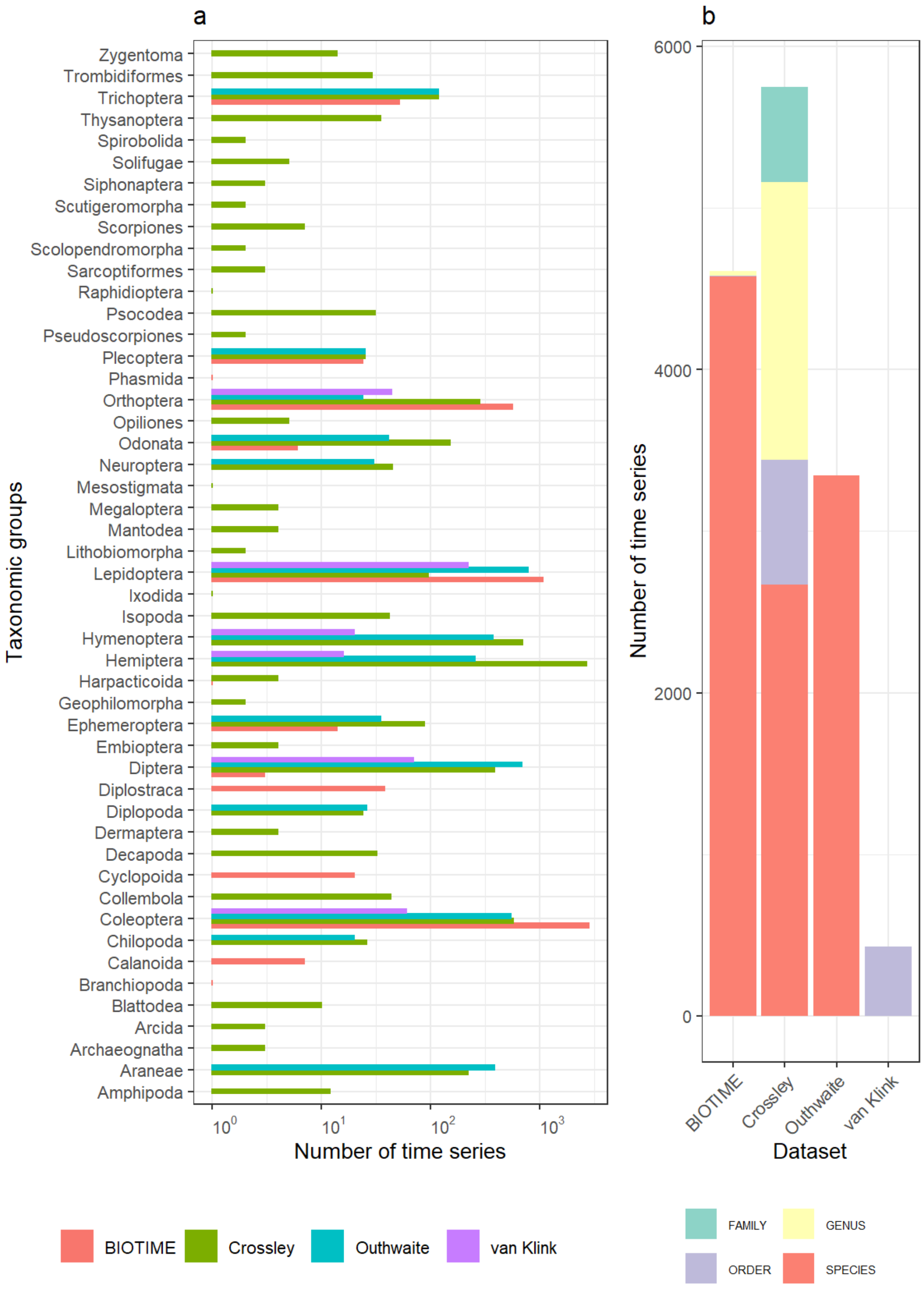

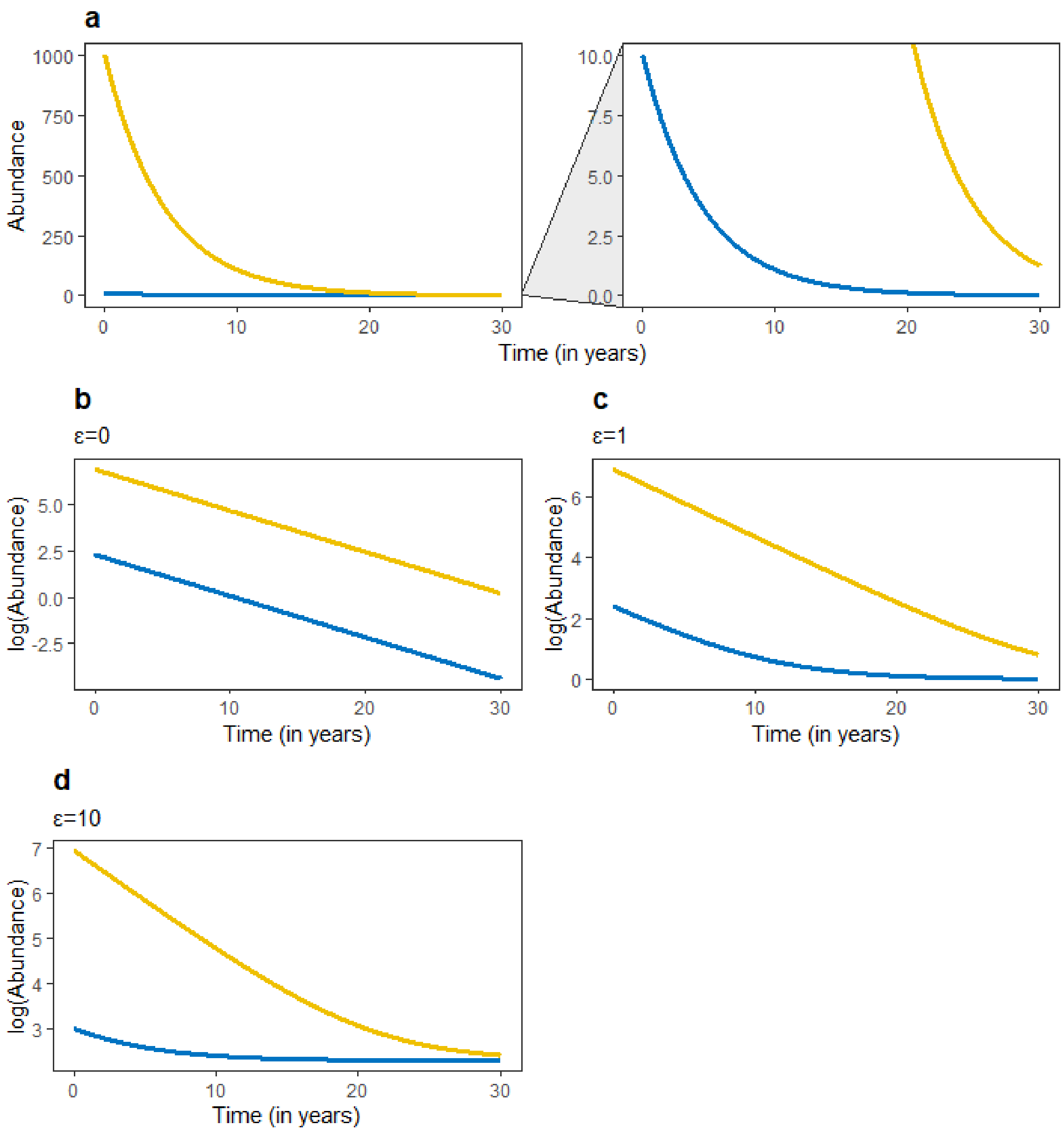


***Fig. S2: Representation of a geometric decline over time.*** *(a) Abundance, measured as number of individuals, against time, for two species declining with a rate of 20% per year (growth rate = 0.8). Zooming on a part on the bottom of the y-axis, we can see that both species decline with the same rate, but since one is rare and the other is common, variations in number of individuals are much larger for the common species. (b,c,d) represents the same dynamics as in a, but applying the transformation log(x+ε) to the abundance, with three values of ε.*

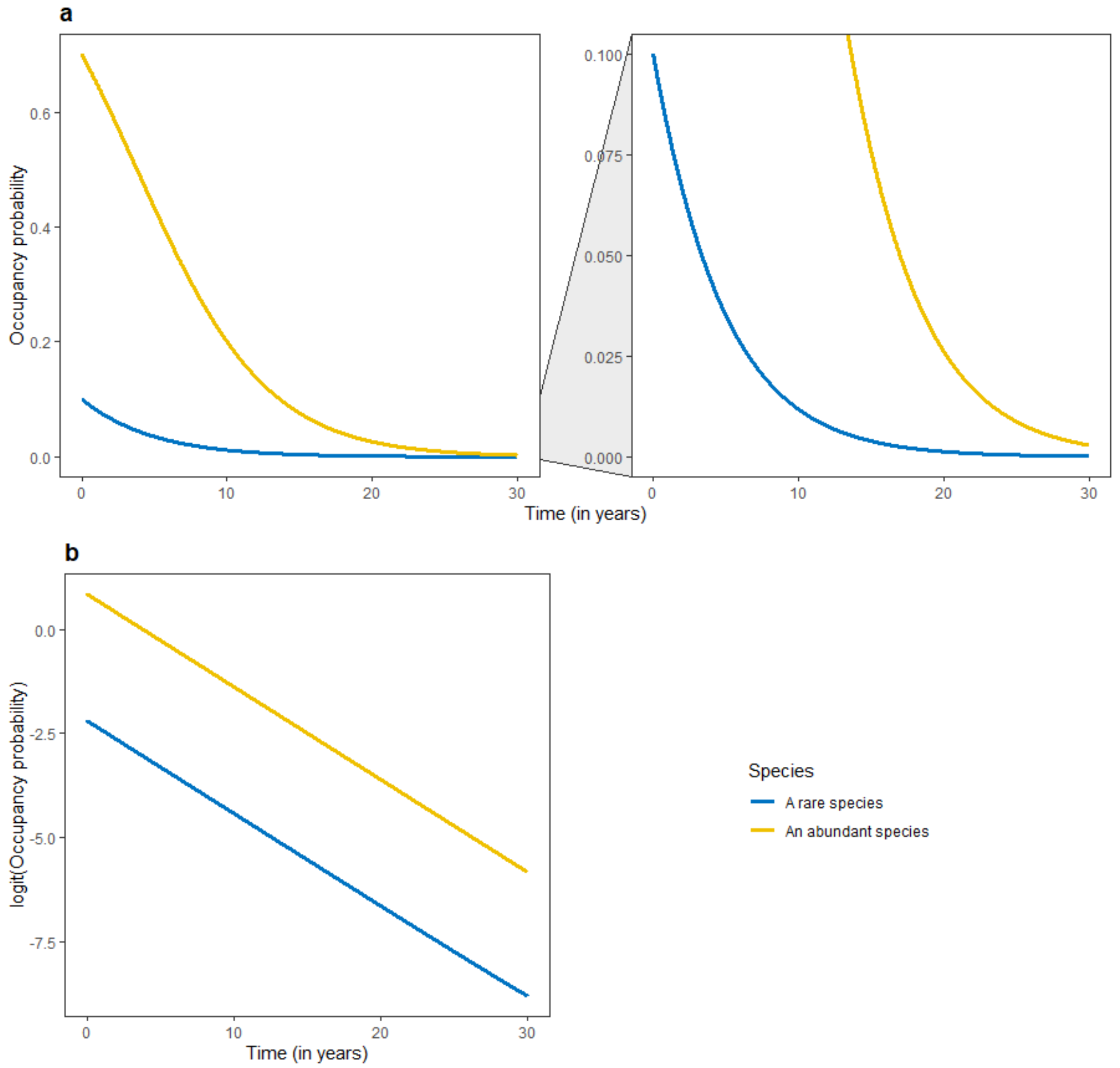


***Fig. S3: Representation of a geometric decline over time.*** *(a) Occupancy probability, measured as the proportion of grid cell occupied by a species on a given area, against time, for two species declining at a growth rate of 0.8, as in figure S2. Zooming on a part on the bottom of the y-axis, we can see that both species decline with the same rate, but since one is rare and the other is common, variations in number of individuals is much larger for the common species. (b) represents the same dynamics as in a, but when applying the transformation logit(x) to the occupancy probability. On the logit scale we can see that declines are linear and identical (same slopes) between both species, and can easily be estimated with a linear model.*

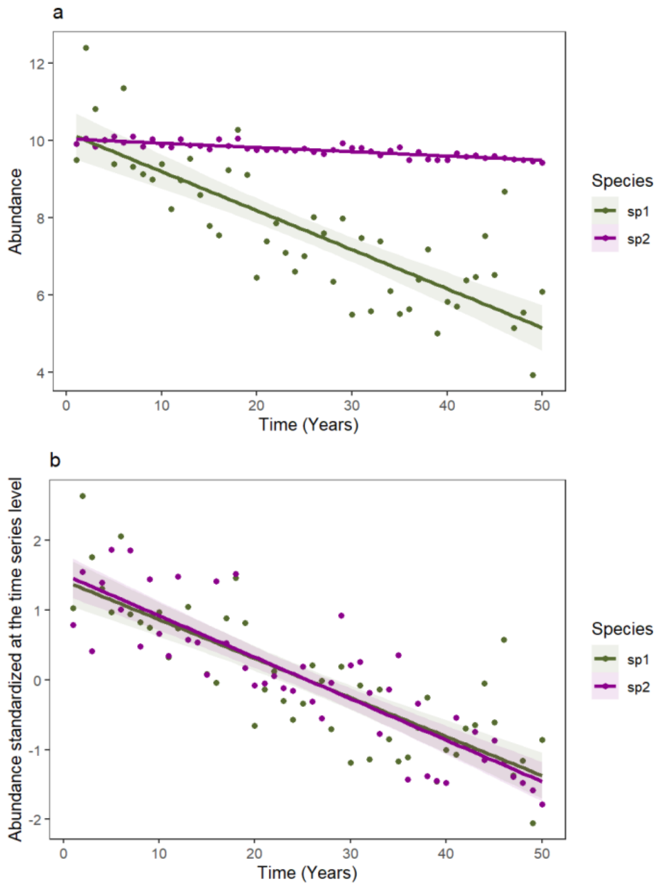


***Fig. S4: Simulated data showing how standardization can affect abundance trends.*** *(a) abundance values over time and associated abundance trends, for one strongly declining species, with high inter-annual variation in abundance (green), and for another species slightly declining, with low inter-annual variation in abundance (purple). (b) shows the same data as in (a) but after standardization of abundance values within each time series (minus mean and then divided by standard deviation). In (b) abundance trends are calculated on standardized data, giving similar trends among species while in fact the green species is declining more than the purple one. Because purple species exhibit low inter-annual variability in abundance, the absolute value of its trend is artefactually increased by the standardization.*

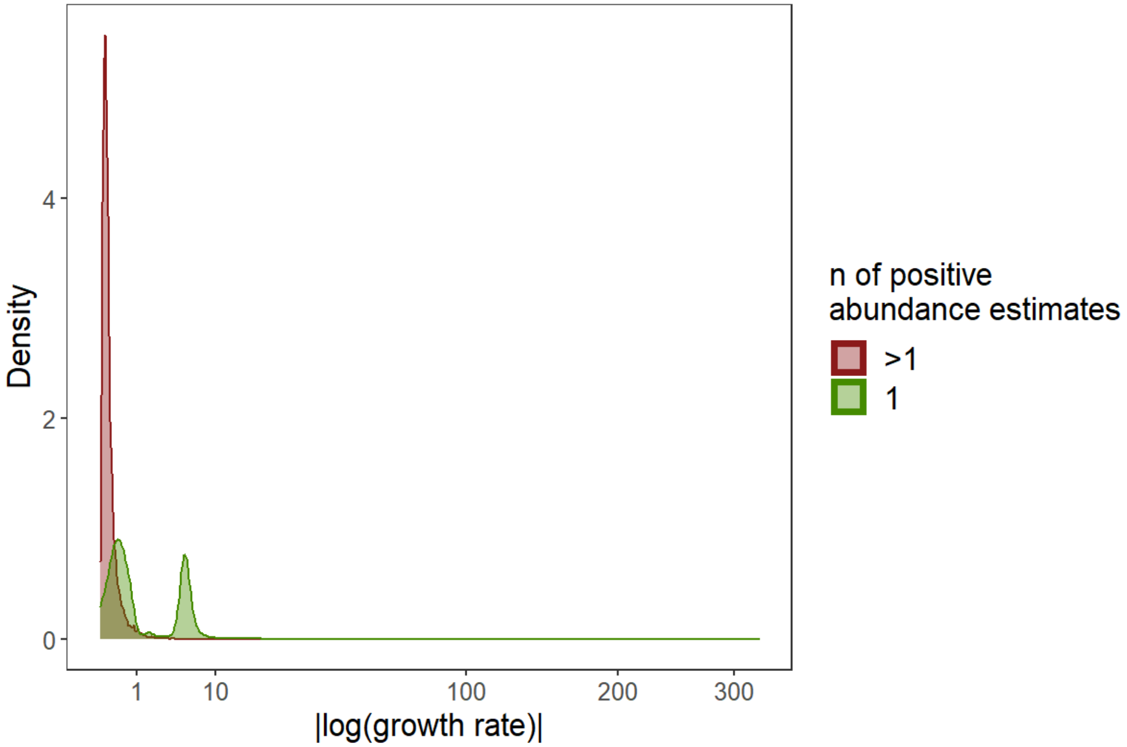


***Fig. S5: Truncated time series with only one non-zero yearly estimate of abundance produce extreme value of abundance trends.*** *Density distribution of the absolute value of abundance trends, log(growth rate), as function of the number of non-zero yearly estimate of abundance contained in the truncated time series (1 vs >1). To preserve readability the x-axis is square-root transformed.*

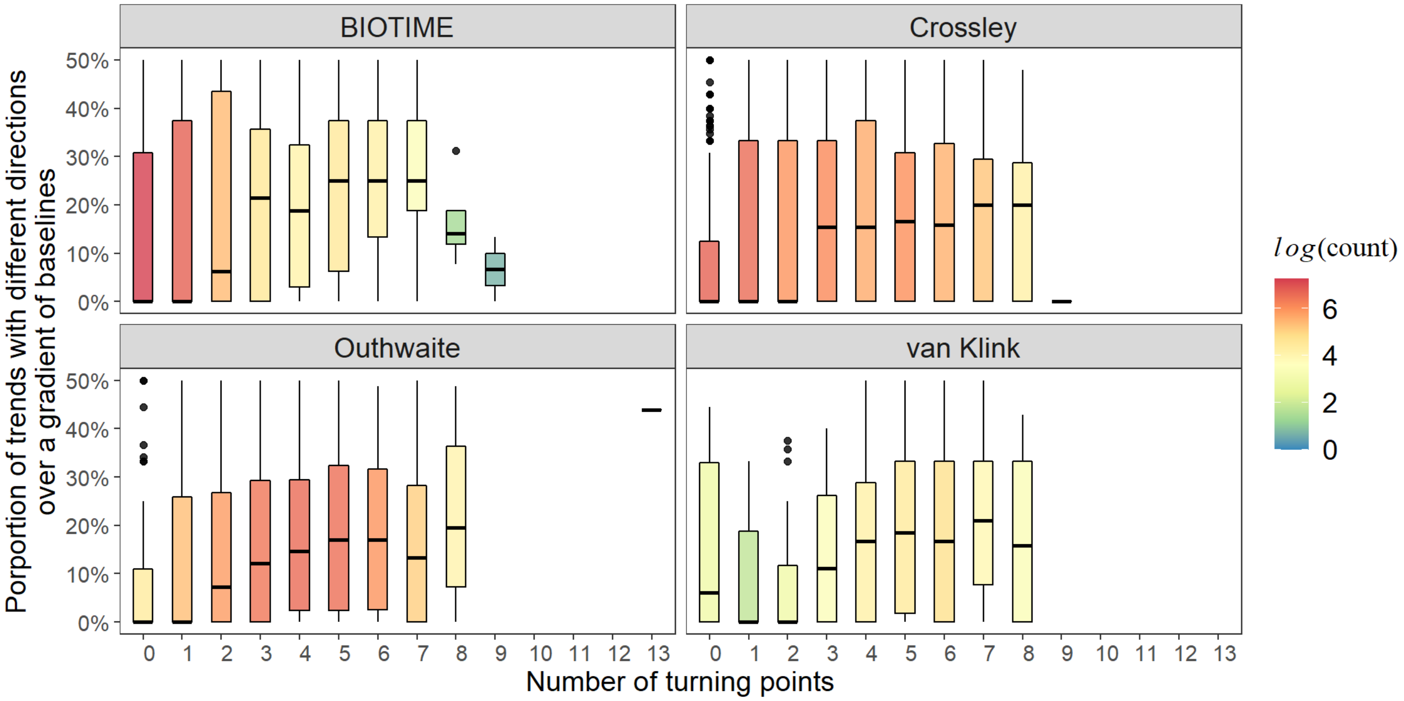


***Fig. S6: Same figure as in the main text (Figure 3b), but for each source dataset.*** *Proportion of abundance trends from truncated time series with different directions (positive vs. negative), as a function of the number of turning points in the corresponding original time series. Boxplots represent minimum and maximum values (bottom and top of vertical lines), first and third quartiles (Q1 and Q3, bottom and top of boxes) and median (thick horizontal lines); colours indicate sample size (number of original time series). Points with values outside of the range [Q1-1.5(Q3-Q1), Q3+1.5(Q3-Q1)] are considered as outliers and represented as full circles.*

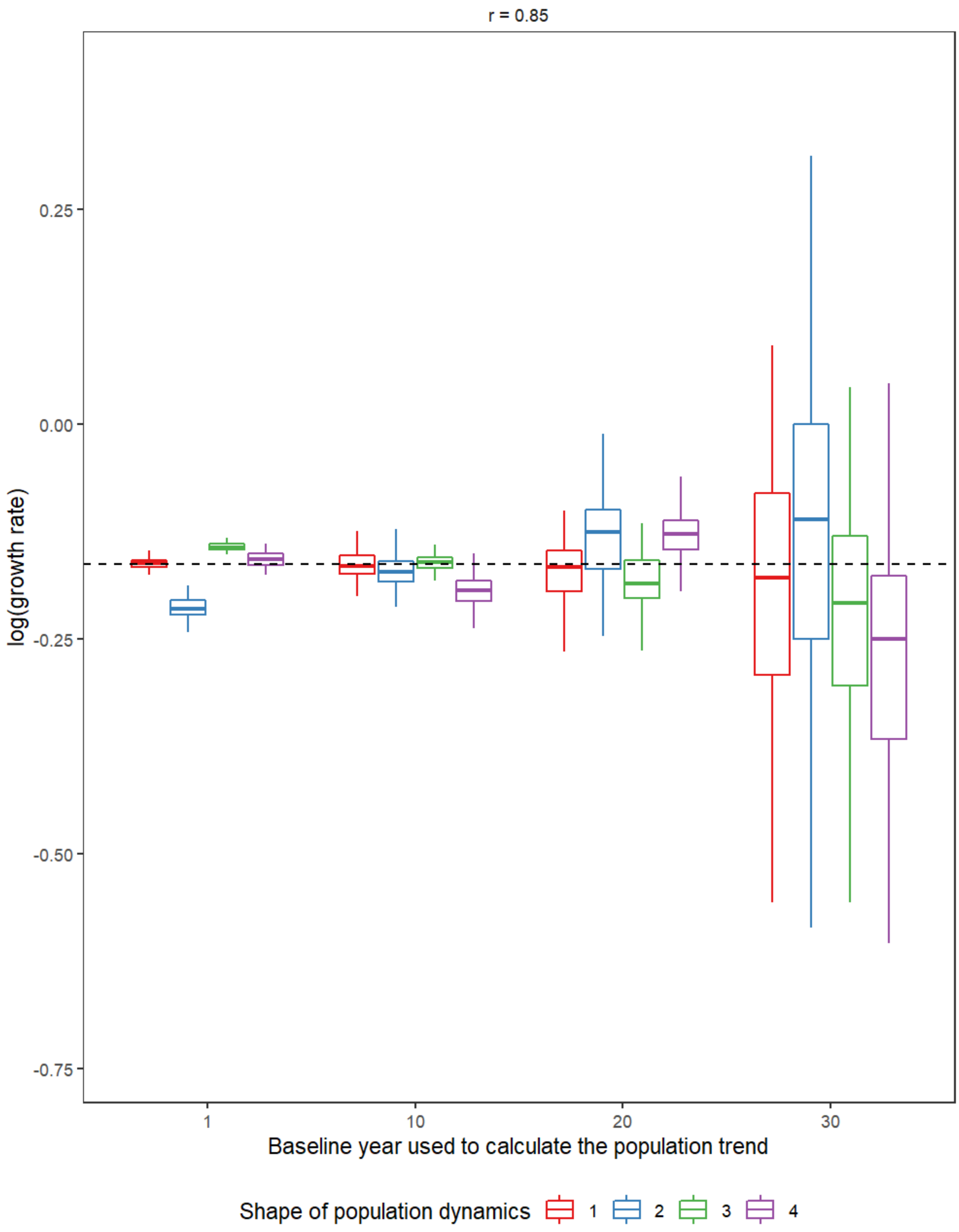


***Fig. S7: Estimated growth rate when a decline of 15%/year is simulated.*** *This is the same plot than in the left panel of Figure 4d, but without the outliers to improve readability of average biases. The dashed horizontal line shows the value of the logarithm of the true (simulated) growth rate.*

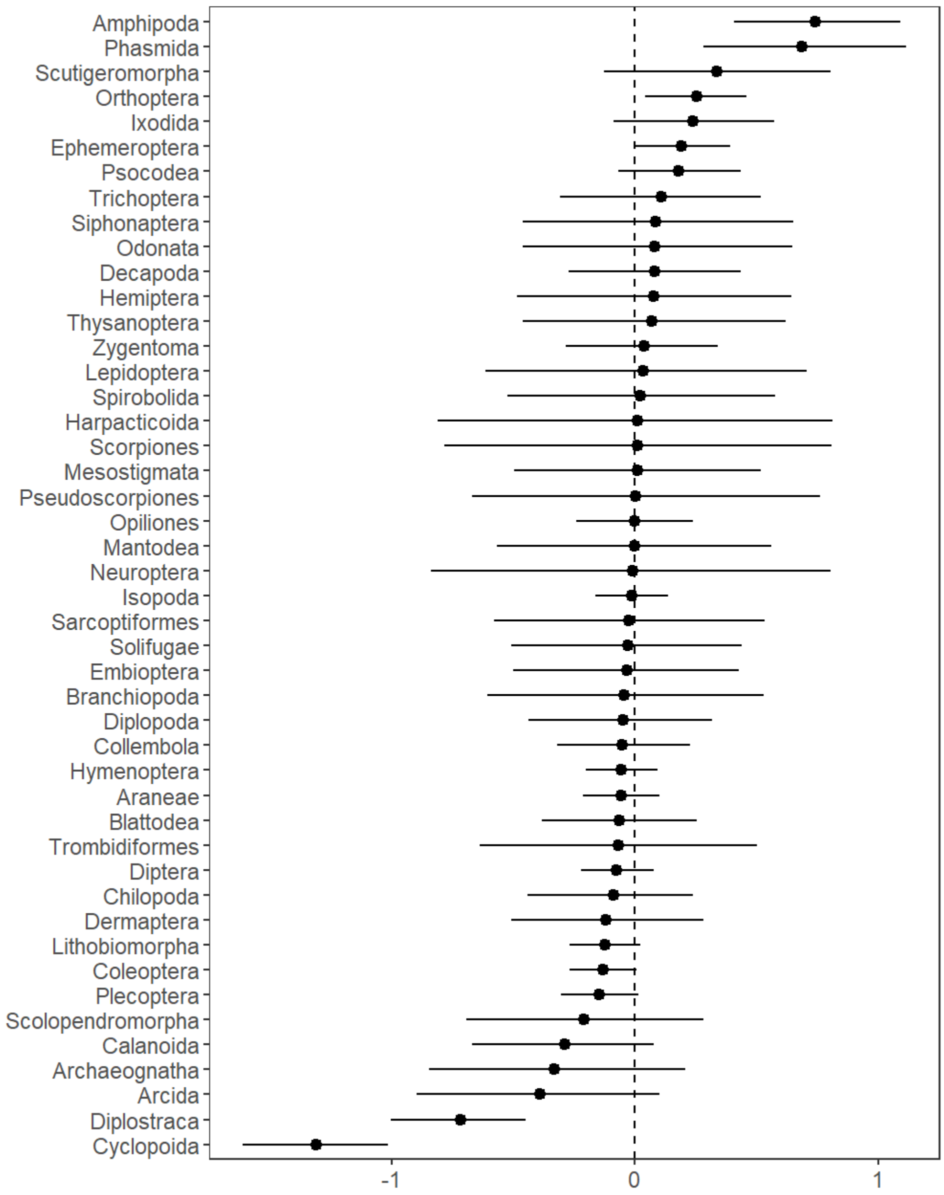


***Fig. S8: Random effects of taxonomic order on abundance trends.*** *The dots show the mean of the posterior distribution while the error bars show the confidence interval at 95% (quantile 2.5% and 97.5%).*

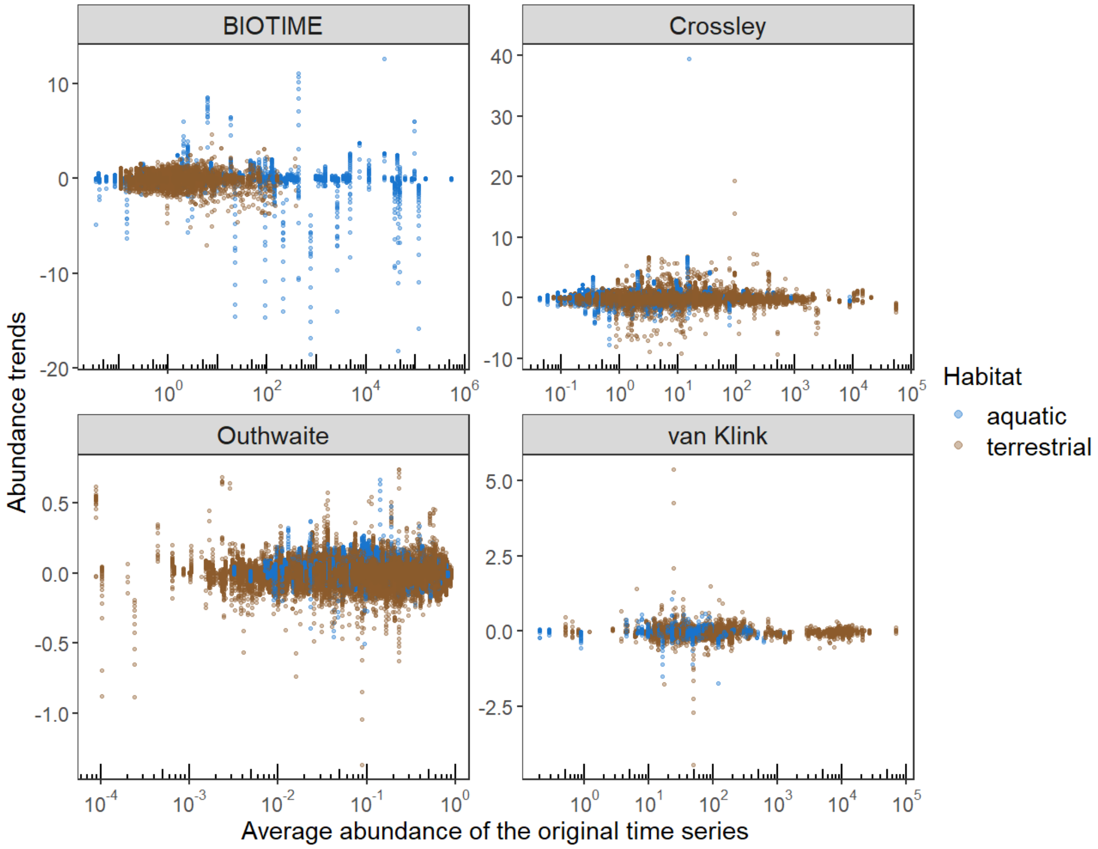


***Fig. S9: Estimated abundance trends of truncated time series as a function of average abundance.*** *Abundance trends (log growth rate) as a function of the average abundance of the original time series (measured on the original abundance scale), per source dataset and habitat.*

**Table S1:** Description of the 4 datasets used.

| Dataset's name | Kind of abundance estimate | Original spatial scope | Spatial scope used here | Taxonomical resolution of time-series | Original temporal coverage | Temporal coverage used here |
|---|---|---|---|---|---|---|
| Outhwaite *et al.* 2019 | Annual occupancy estimate (*i.e.* the proportion of 1km$^2$ grid cells in a region occupied by a species, a proxy for abundance) | Great-Britain or United-Kingdom or region (Wales, England, Scotland, Northern Ireland) levels | Time-series at Great-Britain level | Species level | 1970-2015 | 1970-2015 |
| Crossley *et al.* 2019 | Local annual abundance count | Local time series spread across the USA | Local time series spread across the USA | Species level mostly (cf. Figure S1) | 1943-2019 | 1970-2019 |
| van Klink *et al.* 2020 | Local annual abundance count, aggregated from literature | Local time series spread across the world | Local time series spread across North America and Europe | Order level | 1925-2018 | 1970-2018 |
| BIOTIME database | Local annual abundance count | Local time series spread across the world | Local time series spread across North America and Europe | Species level | 1874-2018 | 1970-2018 |

**Table S2:** Number of original time series across datasets, continents and habitats, at the end of the filtering process.

| Dataset | Continent | aquatic | terrestrial |
|---|---|---|---|
| BIOTIME | Europe | 142 | 3877 |
| BIOTIME | North America | 26 | 562 |
| Crossley | North America | 587 | 5161 |
| Outhwaite | Europe | 265 | 3081 |
| van Klink | Europe | 0 | 101 |
| van Klink | North America | 70 | 258 |
