## Supplementary material for "Controversy over the decline of arthropods: a matter of temporal baseline?": R code

May 13, 2022

### 1 Settings and check that packages are installed:

R version: 4.0.3 Exploitation system: Windows 10

```
knitr::opts_chunk$set(echo = TRUE, fig.cap = " ")
#Before running the script, please set the path to the folder that contains the data.
filepath="C:/Users/Duchenne/Documents/baseline/for_review/data_needed_to_run_script"
setwd(dir=filepath)
library(ggradar)
library(ggplot2)
library(gridExtra)
library(data.table)
library(dplyr)
library(rgbif)
library(ggforce)
library(ggeffects)
library(ggExtra)
library(viridis)
library(lme4)
library(cowplot)
library(scales)
library(car)
library(UKBiodiversity)
library(magick)
library(ggpubr)
library(rjags)
library(R2jags)
library(mcmcplots)
library(MCMCvis)
library(mcmcOutput)
library(ggridges)
library(mgcv)
library(runjags)
library(pastecs)
library(sf)
library(INLA)
library(foreach)
library(doParallel)
library(ggthemes)
```

Here there are all the steps to reproduce analyses from the beginning to the end, with the original data. However, since some steps can take few hours or days to run (especially the estimation of abundance trends), I provided some data table produced by the scripts in the data folder, to allow to reproduce only the final steps (statistics) if needed. The statistical models used in the article, are also provided as Rdata object in the data folder in case reviewers would like to reproduce figure without re-perfomring analyses.

I tried to organize the code chunks in the same order than the methods in the paper, to improve the clarity of the code.

### 2 Data aggregation and cleaning

#### 2.1 Data from Outhwaite et al. (2019)

Load data, aggregate all tables, apply filters and check the taxonomy with the GBIF tool. Then we scaled abundances and put the table in the final format.

```
#Concatenate tables (operation to do once only):
# setwd(dir="C:/Users/duche/Documents/baseline/SUMMARY_TABLES")
# liste=list.files()
# tab=NULL
# for(i in liste){
# temp=fread(i,header=T)
# tab=rbind(tab,temp)
# }
# setwd(dir="C:/Users/duche/Documents/baseline")
# fwrite(tab,"ocucpancy_estimates_byyear_GB.txt")
setwd(dir=filepath)
tab=fread("ocucpancy_estimates_byyear_GB.txt",header=T)

tab=subset(tab,Region=="GB") #take estimates at the Great-Britain level only
tab=subset(tab,Rhat<1.2) # remove estimates which congered badly
#exlude non-arthropods groups:
tab=subset(tab,Group!="Lichens" & Group!="Bryophytes"&
Group!="NonmarineMolluscs")
#merge estimates with a table with information about groups:
tab=merge(tab,major_groups,by=c("Group","Species"))
tab$Habitat="terrestrial"
tab$Habitat[tab$Major_group=="Freshwater species"]="aquatic"

#merge estimates with another table with information about groups:
bidon=fread("group_order_table_UK.txt",header=T)
tab=merge(tab,bidon,by="Group")

bidon=unique(tab[,c("Species","order")])
bidon=subset(bidon,!is.na(order))
for(i in 1:nrow(bidon)){
#look for the given species:
obj=name_backbone(name=bidon[i,"Species"],
```

```
kingdom='Animalia',order=bidon[i,"order"])
if(length(obj$rank)>0){  #if obj is not empty
bidon$rang[i]=obj$rank  #keep the rank of the match
bidon$confi[i]=obj$confidence #confidence in the match
bidon$kingdom[i]=obj$kingdom #kingdom
bidon$canon[i]=obj$canonicalName #latin name according to the GBIF
if(length(obj$order)>0){bidon$order2[i]=obj$order}else{bidon$order2[i]=NA}
#keep the order according to the GBIF
}
print(i)
}

tab=merge(tab,bidon[,c("Species","rang","canon")],by=c("Species"))
tab$Species[tab$order==tab$order2 &
!is.na(tab$canon)]=tab$canon[tab$order==tab$order2 & !is.na(tab$canon)]
tab$Study="Outhwaite"
tab$Continent="Europe"
tab$Rank="SPECIES"

tab=tab %>% group_by(Study,Continent,Group,coverage,order,Species,Habitat)
%>%
  mutate(time_series_id=paste0("OUTH_",cur_group_id())) #create an ID per
time series

tab=tab %>% group_by(time_series_id) %>% mutate(lastyear=max(Year))
#calculate the last year of each time series

tab$abond=scale(logit(tab$Mean,adjust=0.001)) #logit transform abundance
estimates
tab$Period=NA
tab$ABUNDANCE_TYPE="Occupancy"

tab=tab[,c("Study","Continent","Group","coverage","Year","order","Species","H
abitat","Mean","abond",
           "time_series_id","Rank","Period","ABUNDANCE_TYPE","lastyear")]
names(tab)=c("Study","Continent","Datasource","Site","Year","Order","Species"
,"Habitat","Abundance",

"abond","time_series_id","Rank","Period","ABUNDANCE_TYPE","lastyear")

fwrite(tab,"Outhwaite_for_use.txt")
```

#### 2.2 Data from Crossley et al. (2020)

Load data, apply filters and check taxnonmy with the GBIF tool. Then we scaled abundances and put the table in the final format.

```
setwd(dir=filepath)
tab2=fread("External_Database_S1_PerSpecies_Abundance_LTER_annotated.csv",hea
der=T)
tab2[tab2==""]=NA #correct the way to code missing information
```

```
tab2[tab2=="?"]=NA #correct the way to code missing information
tab2$Study="Crossley"
tab2$Continent="North America"

#Taxonomy checks and manual corrections:
tab2$Species.code[tab2$Species.code=="ORTHO" &
tab2$Order=="Diptera"]="Orthocladius"
tab2[grep("C. ",tab2$Species,fixed=T),]
tab2$Species=gsub("C. ","Culex ",tab2$Species,fixed=T)
tab2[grep("Ae. ",tab2$Species,fixed=T),]
tab2$Species=gsub("Ae. ","Aedes ",tab2$Species,fixed=T)
tab2[grep("Oc. ",tab2$Species,fixed=T),]
tab2$Species=gsub("Oc. ","Ochlerotatus ",tab2$Species,fixed=T)
tab2$Species[tab2$Species=='Baetis ("species 3")']="Baetis sp3"

#First checks:
bidon=unique(tab2[,c("Species.code","Order")])
bidon2=subset(bidon,!is.na(Order))
bidon2[bidon2$Species.code %in%
bidon2$Species.code[duplicated(bidon2$Species.code)],]

#2nd check:
bidon=unique(tab2[,c("Species.code","Species")])
obj=bidon[bidon$Species %in% bidon$Species[duplicated(bidon$Species)],]
obj=obj[order(obj$Species),]
obj=subset(obj,!is.na(Species))
tab2$Species.code[tab2$Species %in% obj$Species]=tab2$Species[tab2$Species
%in% obj$Species]

#3rd check:
#information of species names is spread in two columns, compare both to see
which one should be used
bidon=unique(tab2[,c("Species.code","Species")])
bidon=subset(bidon,!is.na(Species))
tab2$sp_length=sapply(tab2$Species,nchar)-
sapply(as.character(tab2$Species.code),nchar)
tab2$Species.code[tab2$sp_length>0 &
!is.na(tab2$Species)]=tab2$Species[tab2$sp_length>0 & !is.na(tab2$Species)]
unique(tab2$Species.code)

#4th checks (check the Order names using the GBIF):
bidon=data.frame(Order=unique(tab2$Order))
for(i in 1:nrow(bidon)){
obj=name_backbone(name=bidon[i,"Order"], kingdom='Animalia') #rechercher
l'espèce voulue
if(length(obj$rank)>0){  #si obj non vide
bidon$rang[i]=obj$rank  #stocker le rang du match (espèce, genre, famille...)
bidon$confi[i]=obj$confidence #confiance ds le match
bidon$kingdom[i]=obj$kingdom
if(length(obj$order)>0){bidon$order[i]=obj$order}else{bidon$order[i]=NA}
}
```

```
}
bidon$order[bidon$Order %in%
c("Thysanura","Megaloptera","Plecoptera","Trichoptera","Diplopoda","Chilopoda
","Collembola")]=bidon$Order[bidon$Order %in%
c("Thysanura","Megaloptera","Plecoptera","Trichoptera","Diplopoda","Chilopoda
","Collembola")]
tab2=merge(tab2,bidon[,c("Order","order")],by="Order")

#5th checks (Order):
bidon=unique(tab2[,c("Species.code","order")])
bidon=subset(bidon,is.na(order))
for(i in 1:nrow(bidon)){
obj=name_backbone(name=gsub(" undet","",bidon[i,"Species.code"],fixed=T),
kingdom='Animalia') #rechercher l'espèce voulue
if(length(obj$rank)>0){  #if obj is not empty
bidon$rang[i]=obj$rank  #keep the rank of the match
bidon$confi[i]=obj$confidence #confidence in the match
bidon$kingdom[i]=obj$kingdom
if(length(obj$order)>0){bidon$order[i]=obj$order}else{bidon$order[i]=NA}
#keep order name
}
}
bidon=subset(bidon,!is.na(order))
#include modification to the data table:
for(i in 1:nrow(bidon)){
tab2$order[tab2$Species.code==bidon$Species.code[i]]=bidon$order[i]
}

#6th checks (taxonomy level and species name homogenization with the GBIF):
tab2$Species.code=gsub('"',"",tab2$Species.code,fixed=T)
bidon=unique(tab2[,c("Species.code","order")])
bidon=subset(bidon,!is.na(order))
for(i in 1:nrow(bidon)){
obj=name_backbone(name=gsub(" undet","",bidon[i,"Species.code"],fixed=T),
kingdom='Animalia') #rechercher l'espèce voulue
if(length(obj$rank)>0){  #if obj is not empty
bidon$rang[i]=obj$rank  #keep the rank of the match
bidon$confi[i]=obj$confidence #confidence in the match
bidon$kingdom[i]=obj$kingdom
bidon$canon[i]=obj$canonicalName
if(length(obj$order)>0){bidon$order2[i]=obj$order}else{bidon$order2[i]=NA}
}
print(i)
}

#correct taxonomy level to consider some groups as order level groups:
bidon$rang[(bidon$order %in%
c("Thysanura","Megaloptera","Plecoptera","Trichoptera","Diplopoda","Chilopoda
","Collembola")) & !(bidon$rang %in%
c("SUBSPECIES","SPECIES","FAMILY","GENUS","ORDER"))]="ORDER"
bidon$order2[bidon$order %in%
```

```
c("Thysanura","Megaloptera","Plecoptera","Trichoptera","Diplopoda","Chilopoda
","Collembola")]=bidon$order[bidon$order %in%
c("Thysanura","Megaloptera","Plecoptera","Trichoptera","Diplopoda","Chilopoda
","Collembola")]

#include modification to the data table:
tab2=merge(tab2,bidon[,c("Species.code","rang","canon")],by=c("Species.code")
)
tab2$Species.code[tab2$order==tab2$order2 &
!is.na(tab2$canon)]=tab2$canon[tab2$order==tab2$order2 & !is.na(tab2$canon)]
tab2$Species.code[tab2$Species.code=="Cicindela viridisticta
arizonensis"]="Cicindela viridisticta"
tab2$rang[tab2$rang=="SUBSPECIES"]="SPECIES"

tab2=tab2 %>%
group_by(Study,Continent,LTER.site,Locale,Species.code,rang,order,Habitat)
%>% mutate(time_series_id=paste0("CROSS_",cur_group_id())) #create an unique
ID per time series

tab2$Species.code[tab2$rang %in%
c("KINGDOM","ORDER","PHYLUM")]=paste0("SP_",tab2$time_series_id[tab2$rang
%in% c("KINGDOM","ORDER","PHYLUM")]) #create a species name when there is not

tab2=subset(tab2,!is.na(Habitat)) #exclude time series for which the habitat
is not defined

tab2=tab2 %>% group_by(time_series_id) %>% mutate(lastyear=max(Year))
#calculate last year of the time series

tab2$abond=scale(log(tab2$Abundance+1)) #log transformation of the abundance
estimates

tab2$Period=NA
tab2$ABUNDANCE_TYPE="Count"

tab2=tab2[,c("Study","Continent","LTER.site","Locale","Year","order","Species
.code","Habitat","Abundance","abond","time_series_id","rang","Period","ABUNDA
NCE_TYPE","lastyear")]
names(tab2)=c("Study","Continent","Datasource","Site","Year","Order","Species
","Habitat","Abundance","abond","time_series_id","Rank","Period","ABUNDANCE_T
YPE","lastyear")
fwrite(tab2,"Crossley_for_use.txt")
```

#### 2.3 Data from Klink et al. (2020)

Load data and apply filters. Then we scaled abundances and put the table in the final format.

```
setwd(dir=filepath)
tab3=fread("aax9931-Van-Klink-SM-CORRECTED-Data-S1.txt",header=T)
taxo=fread("klink_order.txt",header=T)
```

```
tab3=subset(tab3,!is.na(Number) & Unit=="abundance")
amount3=nrow(tab3)
tab3=merge(tab3,taxo,by=c("Datasource_ID"))
tab3$Realm[tab3$Realm=="Freshwater"]="aquatic"
tab3$Realm[tab3$Realm=="Terrestrial"]="terrestrial"
tab3=subset(tab3,order!="Insects" & order!="Invertebrates" &
order!="Arthropod")
tab3=subset(tab3,Continent=="North America" | Continent=="Europe")
tab3$Study="Klink"

#Excluding LTER sites from North America to avoid redundancy with Crossley et
al.
#vec=grep("LTER",tab3$Datasource_name,fixed=T)
#vec2=which(tab3$Continent=="North America")
#vec=vec[vec %in% vec2]
#tab3=tab3[-vec,]
#Excluding BIOTIME data to avoid redundancy with BIOTIME
#tab3=tab3[grep("BioTIME",tab3$Datasource_name,invert=T,fixed=T),]

tab3$Rank="ORDER"
tab3=tab3 %>%
group_by(Study,Continent,Datasource_ID_4INLA,Plot_ID_4INLA,order,Realm) %>%
mutate(time_series_id=paste0("Klink_",cur_group_id())) #create an ID per time
series
tab3$Species=paste0("SP_",tab3$time_series_id)

tab3=tab3 %>% group_by(time_series_id) %>% mutate(lastyear=max(Year,na.rm=T))
#calculate last year of the time series

#tab3=tab3 %>%
group_by(Study,Continent,Datasource_ID_4INLA,Plot_ID_4INLA,Realm,time_series_
id) %>% mutate(abond=scale(log(Number+1)))
tab3$abond=scale(log(tab3$Number+1)) #log transformation of the abundance
estimates
tab3$ABUNDANCE_TYPE="Count"

tab3=tab3[,c("Study","Continent","Datasource_ID_4INLA","Plot_ID_4INLA","Year"
,"order","Species","Realm","Number","abond","time_series_id","Rank","Period_4
INLA","ABUNDANCE_TYPE","lastyear")]
names(tab3)=c("Study","Continent","Datasource","Site","Year","Order","Species
","Habitat","Abundance","abond","time_series_id","Rank","Period","ABUNDANCE_T
YPE","lastyear")
fwrite(tab3,"Klink_for_use.txt")
```

#### 2.4 Data from BIOTIME

```
setwd(dir=filepath)
met=fread("BioTIMEMetadata_24_06_2021.csv",header=T)
met=met[met$REALM %in% c("Terrestrial","Freshwater")]
met=met[grep("invertebrates",met$TAXA,fixed=T),]
met=met[grep("lter",met$WEB_LINK,invert=T),] #remove LTER data
setwd(dir="C:/Users/Duchenne/Documents/baseline/for_review/data_needed_to_run
```

```
_script/continents")
shp=st_read(".","continent")
pts=st_as_sf(met,coords=c("CENT_LONG","CENT_LAT"),crs=st_crs(shp))
pli = st_intersection(pts,shp) # associate data to continent
met=as.data.frame(pli %>% st_drop_geometry())
met=met[met$STUDY_ID %in% pli$STUDY_ID,]
met=met[met$CONTINENT %in% c("North America","Europe"),] #select only data
from North America and from Europe

met$Habitat="aquatic"
met$Habitat[met$REALM=="Terrestrial"]="terrestrial"
met$Study="BIOTIME"
met$Datasource=paste(met$Study,met$STUDY_ID,sep="_")

setwd(dir=filepath)
dat=fread("BioTIMEQuery_24_06_2021.csv",header=T)
dat$Study="BIOTIME"
dat$Datasource=paste(dat$Study,dat$STUDY_ID,sep="_")
a=length(unique(paste0(dat$Datasource,dat$GENUS_SPECIES,dat$PLOT)))
tab4=merge(unique(met[,c("Study","Datasource","Habitat","CONTINENT","ABUNDANC
E_TYPE")]),dat,by=c("Study","Datasource"))
a2=length(unique(paste0(tab4$Datasource,tab4$GENUS_SPECIES,tab4$PLOT)))
tab4=subset(tab4,!is.na(ABUNDANCE_TYPE))

#taxanomic check:
bidon=data.frame(GENUS_SPECIES=unique(tab4$GENUS_SPECIES))
bidon$GENUS_SPECIES=gsub(" undet"," sp",bidon$GENUS_SPECIES,fixed=T)
for(i in 1:nrow(bidon)){
obj=name_backbone(name=bidon[i,"GENUS_SPECIES"], kingdom='Animalia') #look
for the given species
if(length(obj$rank)>0){  #if obj is not empty
bidon$rang[i]=obj$rank  #keep the rank of the match
bidon$confi[i]=obj$confidence #confidence in the match
bidon$kingdom[i]=obj$kingdom #kingdom
bidon$canon[i]=obj$canonicalName #latin name according to the GBIF
if(length(obj$order)>0){bidon$order[i]=obj$order}else{bidon$order[i]=NA}
#keep the order according to the GBIF
}
print(i)
}

list_to_keep=bidon$GENUS_SPECIES[bidon$rang=="GENUS"][grep("
sp",bidon$GENUS_SPECIES[bidon$rang=="GENUS"],invert=T,fixed=T)] #list of
species for which GBIF name should not be trusted
list_to_keep=list_to_keep[!(list_to_keep %in% c("Psinidia","Prays f"))]
bidon$rang[bidon$GENUS_SPECIES %in% list_to_keep]="SPECIES"
#keep some classes and put them at the order level:
bidon$order[bidon$canon=="Turbellaria"]="Turbellaria"
bidon$order[bidon$canon=="Branchiopoda"]="Branchiopoda"
bidon[bidon$GENUS_SPECIES=="Geometrid
sp",c("rang","canon","order")]=c("ORDER","Geometrid sp","Lepidoptera")
```

```
bidon[bidon$GENUS_SPECIES=="Noctuid
sp",c("rang","canon","order")]=c("ORDER","Noctuid sp","Lepidoptera")
#remove non arthropod invertebrates
bidon=subset(bidon,canon!="Oligochaeta")
bidon=subset(bidon,canon!="Turbellaria")
bidon=subset(bidon,order!="Flosculariaceae")
bidon=subset(bidon,order!="Myida")
bidon=subset(bidon,order!="Ploima")
bidon=subset(bidon,order!="Littorinimorpha")
bidon=subset(bidon,order!="Arcida")

tab4=merge(tab4,bidon[,c("GENUS_SPECIES","rang","canon","order")],by=c("GENUS
_SPECIES"))
tab4$GENUS_SPECIES[!(tab4$GENUS_SPECIES %in%
list_to_keep)]=tab4$canon[!(tab4$GENUS_SPECIES %in% list_to_keep)]
tab4$GENUS_SPECIES[!is.na(tab4$canon)]=tab4$canon[!is.na(tab4$canon)]
summary(as.factor(tab4$order))

#generate zero when there is sampling on same locality for the same order
mat=dcast(tab4,Study+CONTINENT+Datasource+PLOT+order+Habitat+YEAR+ABUNDANCE_T
YPE~GENUS_SPECIES,value.var="sum.allrawdata.ABUNDANCE",fun.aggregate =
mean,na.rm=T,fill=0)
mat2=melt(mat,id.vars=c("Study","CONTINENT","Datasource","PLOT","order","Habi
tat","YEAR","ABUNDANCE_TYPE"),variable.name =
"GENUS_SPECIES",value.name="ABUNDANCE")
mat2=mat2 %>%
dplyr::group_by(Study,CONTINENT,Datasource,PLOT,order,GENUS_SPECIES,Habitat,A
BUNDANCE_TYPE) %>%
dplyr::mutate(pres=sum(ABUNDANCE),time_series_id=paste0("BIOTIME_",dplyr::cur
_group_id())) #create a unique Id per time series and calculate the sum of
the abundance per species per locality
mat2=subset(mat2,pres>0) #removing species absent from a locality

tab4=merge(mat2,unique(tab4[,c("GENUS_SPECIES","rang","order")]),by=c("GENUS_
SPECIES","order"))
subset(tab4,Datasource=="BIOTIME_249" & PLOT=="" &
GENUS_SPECIES=="Gravitarmata margarotana")

tab4=tab4 %>% dplyr::group_by(time_series_id) %>%
dplyr::mutate(lastyear=max(YEAR,na.rm=T)) #calculate last year of the time
series

tab4$abond=scale(log(tab4$ABUNDANCE+1))
tab4$Period=NA
tab4$Site=paste(tab4$Study,tab4$Datasource,tab4$PLOT,sep="_")

tab4=tab4[,c("Study","CONTINENT","Datasource","Site","YEAR","order","GENUS_SP
ECIES","Habitat","ABUNDANCE","abond","time_series_id","rang","Period","ABUNDA
NCE_TYPE","lastyear")]
names(tab4)=c("Study","Continent","Datasource","Site","Year","Order","Species
","Habitat","Abundance","abond","time_series_id","Rank","Period","ABUNDANCE_T
```

```
YPE","lastyear")

fwrite(tab4,"BIOTIME_for_use.txt")
```

#### 2.5 Aggregate the 4 datasets together and generating an array of datasets using a sliding baseline

```
setwd(dir=filepath)
tabb=fread("ocucpancy_estimates_byyear_GB.txt",header=T)
tabb=subset(tabb,Region=="GB")
tab=fread("Outhwaite_for_use.txt")
nrow(tab)/nrow(tabb)
tab2b=fread("External_Database_S1_PerSpecies_Abundance_LTER_annotated.csv",he
ader=T)
tab2=fread("Crossley_for_use.txt")
nrow(tab2)/nrow(tab2b)
tab3b=fread("aax9931-Van-Klink-SM-CORRECTED-Data-S1.txt",header=T)
tab3=fread("Klink_for_use.txt")
nrow(tab3)/nrow(tab3b)
summary(tab3$Year)
tab4b=fread("BioTIMEQuery_24_06_2021.csv",header=T)
tab4=fread("BIOTIME_for_use.txt")
nrow(tab4)/nrow(tab4b)
summary(tab4$Year)
unique(tab4$Rank)
tab3$Datasource=as.character(tab3$Datasource)
tab3$Site=as.character(tab3$Site)
tabf=rbind(tab,tab2,tab3,tab4)
tabf$Order[tabf$Order=="Thysanura"]="Zygentoma"
tabf$Rank[tabf$Rank=="KINGDOM"]="ORDER" #Consider remaining groups such as
Diplododa, as order
tabf$Rank[tabf$Rank=="PHYLUM"]="ORDER" #Consider remaining groups such as
Diplododa, as order
a=length(unique(tabf$time_series_id))
tabf=subset(tabf,lastyear>=2005) #exclude time series with a last year
posterior to 2010
a-length(unique(tabf$time_series_id))
a1=nrow(tabf)
tabf=subset(tabf,Year>=1970)
a1-nrow(tabf)
a2=length(unique(tabf$time_series_id))
tabf=tabf %>% group_by(time_series_id) %>% mutate(nbann=length(unique(Year)))
tabf=subset(tabf,nbann>=3)
a2-length(unique(tabf$time_series_id))

unique(tabf$ABUNDANCE_TYPE)

setwd(dir=filepath)
length(unique(tabf$time_series_id))
fwrite(tabf,"compiled_dataset.txt")
```

```
#Creating a palette of datasets (just need to be executed once):
setwd(dir=filepath)
for(i in c(1970:2010)){
bide=subset(bidon2,Year==i) #select first year of time periods
bidon=tabf[tabf$time_series_id %in% bide$time_series_id,] #select time_series
with an occupancy estimate at least for the first year of the period
bidon=subset(bidon,Year>=i) #select time period
bidon$baseline_trunc=i
bidon=bidon %>% group_by(time_series_id) %>%
mutate(nbann=n_distinct(Year),somme=sum(Abundance))
bidon=subset(bidon,nbann>=3 & somme>0)
if(i==1970){tabff=bidon}else{tabff=rbind(tabff,bidon)}
print(paste(i,nrow(tabff)))
}
fwrite(tabff,"dataset_palette.txt")
```

### 3 Abundance trends

#### 3.1 Assessing linearity of originales time series

```
setwd(dir=filepath)

tab=fread("compiled_dataset.txt")
tab$Abundance_trans=tab$Abundance
tab$Abundance_trans[tab$Study=="Outhwaite"]=logit(tab$Abundance[tab$Study=="O
uthwaite"],adjust=1e-14)
tab=subset(tab,!is.na(Abundance_trans))

liste=as.data.frame(unique(tab,
by=c("time_series_id","Study","Continent","Datasource","Site","Order","Specie
s","Habitat","Rank")))
liste$edf=NA
liste$nbann=NA
liste$baseline=NA
liste$nturns=NA
liste$r.sq=NA
liste$abond_moy=NA

for(i in unique(liste$time_series_id)){
bidon=subset(tab, time_series_id==i)
bidon=subset(bidon,!is.na(Abundance_trans))
bidon=bidon[order(bidon$Year),]
if(i=="CROSS_1378"){bidon$Abundance_trans=abs(bidon$Abundance_trans)}
    if(bidon$ABUNDANCE_TYPE %in%
c("Occupancy","Presence/Absences")){famille="gaussian"}
    if(bidon$ABUNDANCE_TYPE %in% c("Count")){famille="poisson"}
    if(bidon$ABUNDANCE_TYPE %in% c("Density")){famille="gaussian(log)"}
if(sum(abs(bidon$Abundance))>0){
if(nrow(bidon)>2){
```

```
if(length(unique(bidon$Period))>1){
model=gam(Abundance_trans~s(Year,k=min(c(length(unique(bidon$Year))-
1,10)))+s(Period, bs = 're'), data=bidon,family=famille)
}else{
model=gam(Abundance_trans~s(Year,k=min(c(length(unique(bidon$Year)),10))),
data=bidon,family=famille)
}
liste$edf[liste$time_series_id==i]=summary(model)$edf[1]
liste$nturns[liste$time_series_id==i]=turnpoints(c(predict(model,terms="s(Yea
r)")))$nturns
liste$r.sq[liste$time_series_id==i]=summary(model)$r.sq
}}
liste$nbann[liste$time_series_id==i]=length(unique(bidon$Year))
liste$baseline[liste$time_series_id==i]=min(bidon$Year)
liste$abond_moy[liste$time_series_id==i]=mean(bidon$Abundance,na.rm=T)

}

fwrite(liste,"linearity_time_series.txt")
```

#### 3.2 Simulations

```
library(doParallel)
library(foreach)
cl<-makeCluster(6)
registerDoParallel(cl)

ntrends=400
years=0:41
middle=max(years)/2
maxi=max(years)
growth_vec=seq(-0.15,0,0.05)
baseline_vec=seq(0,30,10)
startvalue=100

#3 shapes of function
func1=function(x,y,z){z*(1+y)^x}

func2=function(x,y,z){
y2=z*(1+y)^x
return(exp(((1/20*x^2)-(41/20*x))/20+log(y2)))}

func3=function(x,y,z){
y2=z*(1+y)^x
return(exp(((-1/20*x^2)+(41/20*x))/40+log(y2)))}

func4=function(x,y,z){
y2=z*(1+y)^x
return(exp((-cos(x/2.956305)+sin(x/4))/4+log(y2)))}

vec=c(func1,func2,func3,func4)
```

```
growth=0
comb <- function(...) {
  mapply('rbind', ..., SIMPLIFY=FALSE)
}
res=foreach(i=1:ntrends,.combine=comb)%dopar%{
library(INLA)
resultat=NULL
resultat2=NULL
years=0:41
startvalue=100
maxi=max(years)
nfun=ceiling(i/(ntrends/4))#sample(1:3,1)

for(growth in growth_vec){
bidon=data.frame(Year=years)
bidon$value_moy=mapply(vec[[nfun]],years,growth,startvalue)
bidon$value=mapply(rpois,n=1,lambda=bidon$value)
bidon$Year=bidon$Year+1
for(base in baseline_vec){
b=subset(bidon,Year>=base)
b$sp=i
b$growth=growth
b$func=nfun
b$baseline=ceiling(base)
resultat=rbind(b,resultat)
b$Year2=b$Year-mean(b$Year)
model=inla(value~Year2+f(Year, model = 'rw1'),
data=b,family="poisson",control.family=list(link="log"),num.threads=1)
b2=model$summary.fixed["Year2",]
b2$sp=i
b2$growth=growth
b2$func=nfun
b2$baseline=base
resultat2=rbind(b2,resultat2)
}
}
return(list(resultat,resultat2))
}
filepath="C:/Users/Duchenne/Documents/baseline/for_review/data_needed_to_run_
script"
setwd(dir=filepath)
save(res, file="simues.RData",version = 2)
stopCluster(cl)
closeCluster(cl)
```

#### 3.3 Estimating abundance trends

```
#Execute on a cluster
library(data.table)
```

```r
library(foreach)
library(doParallel)
# setup parallel backend to use all available CPUs
ncpus <- as.numeric(6)
registerDoParallel(ncpus)

tabf=fread("dataset_palette.txt")
#remove NA:
tabf=subset(tabf,!is.na(Abundance))
#prepare the table
liste=unique(tabf,
by=c("baseline_trunc","time_series_id","Study","Continent","Datasource","Site
","Order","Species","Habitat","Rank"))[,c("baseline_trunc","time_series_id","
Study","Continent","Datasource","Site","Order","Species","Habitat","Rank")]

part=data.frame(time_series_id=unique(liste$time_series_id),part=cut(1:length
(unique(liste$time_series_id)),10))
liste=merge(liste,part,by="time_series_id")
part_vec=unique(liste$part)

for(j in 1:2){
  liste2=subset(liste,part==part_vec[j])
  dim(liste2)

  resultat=NULL
  resultat=foreach(i=unique(liste2$time_series_id),.combine=rbind)%dopar%{
    library(data.table)
    library(INLA)
    a="avec autocorrelation"
    bidon=subset(tabf, time_series_id==i)
    bidon$group=1
    if(i=="CROSS_1378"){bidon$Abundance=abs(bidon$Abundance)}
    bidon=bidon[order(bidon$baseline_trunc,bidon$Year)]
    bidon$baseline_trunc=as.factor(bidon$baseline_trunc)
    bidon$Abundance_round=round(bidon$Abundance)
    res=subset(liste2, time_series_id==i)
    if(bidon$ABUNDANCE_TYPE %in% c("Occupancy","Presence/Absences")){
    famille="xbinomial"
    link="logit"}
    if(bidon$ABUNDANCE_TYPE %in% c("Count")){
      famille="xpoisson"
      link="log"}
    if(bidon$ABUNDANCE_TYPE %in% c("Density")){
      famille="gaussian"
      link="log"}
    res=NULL
    for(jj in unique(bidon$baseline_trunc)){
      bidon2=subset(bidon,baseline_trunc==jj)
      bidon2$Year2=bidon2$Year-min(bidon2$Year)
      model=tryCatch({
```

```
      if(length(unique(bidon$Period))>1){
        inla(Abundance~Year2+f(Year, model = 'rw1')+f(Period, model = 'iid'),
data=bidon2,family=famille,control.family=list(link=link),num.threads=1,blas.
num.threads=1,verbose=F)
      }else{
        inla(Abundance~Year2+f(Year, model = 'rw1'),
data=bidon2,family=famille,control.family=list(link=link),num.threads=1,blas.
num.threads=1,verbose=F)
      }}, error=function(cond) {
        message("fail")
        # Choose a return value in case of error
        return("sans autocorrelation")
      })

      if(class(model)!="inla"){
        if(length(unique(bidon$Period))>1){
          model=inla(Abundance~Year2+f(Period, model = 'iid'),
data=bidon2,family=famille,control.family=list(link=link),num.threads=1,blas.
num.threads=1,verbose=F)
        }else{
          model=inla(Abundance~Year2,
data=bidon2,family=famille,control.family=list(link=link),num.threads=1,blas.
num.threads=1,verbose=F)
        }
      a="sans autocorrelation"
      }

      eff=model$summary.fixed["Year2",]
      eff$baseline_trunc=as.numeric(as.character(jj))
      eff$nbann_pos=length(unique(bidon2$Year[bidon2$Abundance>0]))
      eff$nbann=length(unique(bidon2$Year))
      eff$type=a
      res=rbind(res,eff)

    }

    res=cbind(res,subset(liste2, time_series_id==i))

    return(as.data.frame(res))
  }

  fwrite(as.data.table(resultat),paste0("INLA_population_trends_",j,".txt"))
}

stopCluster(cl)
```

The script estimating abundance trends (above) produce results in ten parts to avoid heavy export from the cluster. So now we aggregate them together:

```
setwd(dir=filepath)
for(j in 1:10){
tabtrt=fread(paste0("INLA_population_trends_",j,".txt"))
tabtrt=tabtrt[,-13]
if(j==1){tabtr=tabtrt}else{tabtr=rbind(tabtr,tabtrt)}
}
tabtr=subset(tabtr,!is.na(sd))
tabtr$Study[tabtr$Study=="Klink"]="van Klink"

tabtr$trend=tabtr$mean

fwrite(tabtr,"tab_trend.txt")
```

### 4 Evaluating the effect of baseline year in empirical abundance trends

#### 4.1 Prepare the data for JAGS model

```
setwd(dir=filepath)
tabtr=fread("tab_trend.txt")
tabtr$dat_site=paste(tabtr$Datasource,tabtr$Site)
# build orthogonal polynomial variable of degree 3 using poly:
pol=as.data.frame(poly(tabtr$baseline_trunc,3))
pol=round(pol,digits=11)
names(pol)[1:3]=paste0("p",names(pol)[1:3])
tabtr=cbind(tabtr,pol)
#convert to numeric indices all factors (more easy to handle in JAGS then)
tabtr$Habitat_num=as.numeric(factor(tabtr$Habitat,levels=sort(unique(tabtr$Ha
bitat))))
tabtr$Continent_num=as.numeric(factor(tabtr$Continent,levels=sort(unique(tabt
r$Continent))))
tabtr$Study_num=as.numeric(factor(tabtr$Study,levels=sort(unique(tabtr$Study)
)))
tabtr$Order_num=as.numeric(factor(tabtr$Order,levels=sort(unique(tabtr$Order)
)))
tabtr$Datasource_num=as.numeric(factor(tabtr$Datasource,levels=sort(unique(ta
btr$Datasource))))
tabtr$dat_site_num=as.numeric(factor(tabtr$dat_site,levels=sort(unique(tabtr$
dat_site))))
tabtr$time_series_id_num=as.numeric(factor(tabtr$time_series_id,levels=sort(u
nique(tabtr$time_series_id))))
tabtr$baseline_trunc=as.numeric(tabtr$baseline_trunc)

#excluding time series with unique positive abundance values:
tabtr_withoutunique=subset(tabtr,nbann_pos>1 & nbann>=3)

#excluding time series with unique positive abundance values:
#create the list of data for JAGS model
Dat<-list(Y=tabtr_withoutunique$trend,
Continents=tabtr_withoutunique$Continent_num,
```

```
Habitats=tabtr_withoutunique$Habitat_num,
Studies=tabtr_withoutunique$Study_num,
Orders=tabtr_withoutunique$Order_num,
Datasources=tabtr_withoutunique$Datasource_num,
dat_sites=tabtr_withoutunique$dat_site_num,
time_series_ids=tabtr_withoutunique$time_series_id_num,
p1=tabtr_withoutunique$p1,
p2=tabtr_withoutunique$p2,
p3=tabtr_withoutunique$p3,
trend_error=tabtr$sd,
unique_Habitat=1:max(tabtr_withoutunique$Habitat_num),
unique_Continent=1:max(tabtr_withoutunique$Continent_num),
unique_Study=1:max(tabtr_withoutunique$Study_num),
unique_Order=1:max(tabtr_withoutunique$Order_num),
unique_Datasource=1:max(tabtr_withoutunique$Datasource_num),
unique_dat_site=1:max(tabtr_withoutunique$dat_site_num),
unique_time_series_id=1:max(tabtr_withoutunique$time_series_id_num),
nbanns=tabtr_withoutunique$nbann,
numberoftrends=nrow(tabtr_withoutunique),
baseline_year=tabtr_withoutunique$baseline_trunc-
min(tabtr_withoutunique$baseline_trunc)+1,
Nbaseline=length(unique(tabtr_withoutunique$baseline_trunc)))

#export it
save(Dat, file="Data_for_hgml_withoutunique.RData",version = 2)
```

#### 4.2 JAGS model

```
model_string<-"model{

for (x in 1:numberoftrends){
#Mean model
fixed[x] <- intercept[Continents[x],Habitats[x]]+Study[Studies[x]]

random[x] <-
baseline[Continents[x],Habitats[x],baseline_year[x]]+Order[Orders[x]]+
#Order random effet
Datasource[Datasources[x]]+  #Datasource random effet
dat_site[dat_sites[x]]+  #site random effet
time_series_id[time_series_ids[x]] #time-series random effet

trends[x]=fixed[x]+random[x]

Y[x] ~ dnorm(trends[x], sigma[x])
sigma[x] <- 1/(var.e[x])

#Dispersion model
log(var.e[x])=intercept_dispersion[Studies[x]]+error_effect*trend_error[x]+nb
ann[Studies[x]]*nbanns[x]+nbann2[Studies[x]]*(nbanns[x]^2)
```

```
#residuals:
E[x] <- (Y[x] - trends[x])

}

# PRIORS

#Mean model priors:

#Order
for (i in unique_Order){
Order[i] ~ dnorm(0, order_tau)
}
order_tau <- 1/(sd.o * sd.o)
sd.o ~ dt(0, 1, 1)T(0,)

#datasource
for (i in unique_Datasource){
Datasource[i] ~ dnorm(0, data_tau)
}
data_tau <- 1/(sd.a * sd.a)
sd.a ~ dt(0, 1, 1)T(0,)

#dat_site
for (i in unique_dat_site){
dat_site[i] ~ dnorm(0, dat_site_tau)
}
dat_site_tau <- 1/(sd.s * sd.s)
sd.s ~ dt(0, 1, 1)T(0,)

#time_series
for (i in unique_time_series_id){
time_series_id[i] ~ dnorm(0, ts_tau)
}
ts_tau <- 1/(sd.ts * sd.ts)
sd.ts ~ dt(0, 1, 1)T(0,)

for (i in unique_Continent){
for (j in unique_Habitat){
intercept[i,j] ~ dnorm(0,0.0001)
}
}

baseline[1,1,1]=0
baseline[1,2,1]=0
baseline[2,1,1]=0
baseline[2,2,1]=0
for (i in unique_Continent){
for (j in unique_Habitat){
```

```
for(k in 2:Nbaseline){
baseline[i,j,k] ~ dnorm(baseline[i,j,(k-1)],tau.baseline)
}
}
}
tau.baseline <- 1/(sd.baseline * sd.baseline)
sd.baseline ~ dt(0, 1, 1)T(0,)

Study[1]<-0
for (i in 2:4){
Study[i]~dnorm(0,0.0001)
}

#Variance Model priors:

for (i in 1:4){
intercept_dispersion[i]~dnorm(0,0.0001)
nbann[i]~dnorm(0,0.0001)
nbann2[i]~dnorm(0,0.0001)
}

error_effect ~ dnorm(0,0.01)T(0,)

####R squared
rsqua<-pow(sd(trends),2)/(pow(sd(trends),2)+pow(sd(E),2))
rsqua_fixed<-pow(sd(fixed),2)/(pow(sd(trends),2)+pow(sd(E),2))
sd.resi<-sd(E)
sd.fixed=sd(fixed)
rsqua2<-pow(sd(trends),2)/(pow(sd(Y),2))
rsqua_fixed2<-pow(sd(fixed),2)/pow(sd(Y),2)

}
"

setwd(dir=filepath)
writeLines(model_string,con="modelcat_hglm.txt")
```

#### 4.3 Run JAGS model

```
library(runjags)
library(rjags)
library(R2jags)
library(mcmcplots)
library(MCMCvis)

# Collect command arguments
args <- commandArgs(trailingOnly = TRUE)
args_contents <- strsplit(args, ' ')
# Get first argument
```

```
j <- as.numeric(args_contents[[1]])
load("Data_for_hgml_withoutunique.RData")

set.seed(j)
#Parameters to track
ParsStage <-
c("intercept","Study","Order","Datasource","dat_site","time_series_id","inter
cept_dispersion","nbann","nbann2",
"sd.a","sd.s","sd.o","sd.ts","sd.resi","sd.fixed","sd.baseline","rsqua","rsqu
a_fixed","rsqua2","rsqua_fixed2","baseline","error_effect")

Inits <- function() {list()}

# RUN MODEL
t1=Sys.time()
model=jags(data=Dat,inits=Inits,parameters.to.save=ParsStage,model.file="mode
lcat_hglm.txt",n.thin=3,n.iter=60000,n.burnin=50000,n.chains=1)
t2=Sys.time()

t2-t1

save(model, file=paste0("model_hglm_withoutunique_",j,".RData"),version =
2)#record_mod6
```

#### 4.4 Combine chains

```
setwd(dir=filepath)

load("model_hglm_withoutunique_1.RData")
model1=model
load("model_hglm_withoutunique_2.RData")
model2=model
load("model_hglm_withoutunique_3.RData")
model3=model
obj1=as.mcmc(model1)
obj2=as.mcmc(model2)
obj3=as.mcmc(model3)
#combine chains and get a summary:
mco <- mcmcOutput(mcmc.list(obj1,obj2,obj3))
bidon=summary(mco)
bidon$varia=rownames(bidon)
fwrite(bidon,"summary_model_withoutunique2.txt")

#collapse chains:
obj=combine.mcmc(list(obj1,obj2,obj3))
save(obj,file="modelchains_withoutunique2.RData",version = 2)
```

### 5 RESULTS

#### 5.1 linearity, figure 3

```
setwd(dir=filepath)
liste=fread("linearity_time_series.txt")

liste$edf2=round(liste$edf,digits=1)
liste$group=paste0((floor(liste$nbann/10)+1)*10-
10,"<n<",(floor(liste$nbann/10)+1)*10)
liste$group[liste$group=="0<n<10"]=" n<10"

p=ggplot(liste, aes(x=nturns,fill=as.factor(group),color=as.factor(group))) +
geom_histogram(aes(y=..density..),bins=14,position=position_dodge2(preserve="
single"),size=0.8,col=NA)+
scale_fill_viridis(discrete
=T,option="E",direction=1)+scale_color_viridis(discrete
=T,option="E",direction=1)+
theme_bw()+ylab("\nDensity\n")+xlab("Number of turning points\n")+
theme(plot.title =
element_text(face="bold",size=14),axis.text=element_text(size=11),axis.title=
element_text(size=14),legend.text=element_text(size=14),legend.title=element_
text(size=14),strip.text= element_text(size=14),
panel.grid=element_blank(),panel.background =
element_rect(fill="white"))+scale_x_continuous(breaks=0:13,labels=function(x)
paste0("  ",x))+labs(fill="number of years",col="number of
years")+ggtitle("a")+coord_cartesian(expand=F)

tabtr=fread("tab_trend.txt")
tabtr=subset(tabtr,nbann_pos>1 & nbann>=3)
bidon=tabtr %>% dplyr::group_by(time_series_id) %>%
dplyr::summarise(ntrends=length(unique(baseline_trunc)),
npos=length(unique(trend[trend<0])),moy=mean(trend,na.rm=T),variance=var(tren
d,na.rm=T))
liste=merge(liste,bidon,by="time_series_id")
liste$val=-1*abs(liste$npos/liste$ntrends-0.5)+0.5

liste=liste %>% dplyr::group_by(nturns) %>%
dplyr::mutate(nbtimseries=length(time_series_id))
p2=ggplot(liste,aes(x=factor(nturns,levels=as.character(0:13)),
y=val,fill=log(nbtimseries))) +
scale_y_continuous(labels=function(x) paste0(x*100, "%"))+
theme_bw()+ylab("Porportion of trends with different directions\nover a
gradient of baselines")+xlab("Number of turning points")+
theme(plot.title =
element_text(face="bold",size=14),axis.text=element_text(size=11),axis.title=
element_text(size=14),legend.text=element_text(size=14),strip.text=
element_text(size=14),panel.grid=element_blank(),legend.position="right",,leg
end.title=element_text(family="serif",size=14))+ggtitle("b")+
scale_x_discrete(breaks=as.character(0:13),labels=function(x) paste0("
```

```
",x),drop=F)+scale_fill_distiller(palette =
"Spectral")+geom_boxplot(width=0.5,
color="black",alpha=0.8)+labs(fill=expression(paste(italic(log),"(count)")))

#leg1 <- get_legend(p)
#p=p+theme(legend.position="none")

setwd(dir=filepath)
pdf("figure_3.pdf",width=8,height=8)
#gridExtra::grid.arrange(p, leg1,p2,ncol=2,widths=c(5,1),heights=c(1.2,1))
plot_grid(p,p2,ncol=1,align="v",rel_heights=c(1.2,1))
dev.off();

liste$Study[liste$Study=="Klink"]="van Klink"
liste=liste %>% dplyr::group_by(nturns,Study) %>%
dplyr::mutate(nbtimseries=length(time_series_id))
p3=ggplot(liste,aes(x=factor(nturns,levels=as.character(0:13)),
y=val,fill=log(nbtimseries))) +
scale_y_continuous(labels=function(x) paste0(x*100, "%"))+
theme_bw()+ylab("Porportion of trends with different directions\nover a
gradient of baselines")+xlab("Number of turning points")+
theme(plot.title =
element_text(face="bold",size=14),axis.text=element_text(size=11),axis.title=
element_text(size=14),legend.text=element_text(size=14),strip.text=
element_text(size=14),panel.grid=element_blank(),legend.position="right",lege
nd.title=element_text(family="serif",size=14))+
scale_x_discrete(breaks=as.character(0:13),labels=function(x) paste0("
",x),drop=F)+scale_fill_distiller(palette =
"Spectral")+geom_boxplot(width=0.5,
color="black",alpha=0.8)+labs(fill=expression(paste(italic(log),"(count)")))+
facet_wrap(~Study)

setwd(dir=filepath)
png("figure_S6.png",width=2000,height=1000,res=200)
#gridExtra::grid.arrange(p, leg1,p2,ncol=2,widths=c(5,1),heights=c(1.2,1))
p3
dev.off();
```

#### 5.2 Simulations, Figure 4

```
setwd(dir=filepath)
load("simues.RData")
resultat=res[[1]]
resultat2=res[[2]]
resultat$growthlabel=factor(paste0("r = ",1+resultat$growth),c("r = 1","r =
0.95","r = 0.9","r = 0.85"))
resultat2$growthlabel=factor(paste0("r = ",1+resultat2$growth),c("r = 1","r =
0.95","r = 0.9","r = 0.85"))
resultat2$baseline[resultat2$baseline==0]=1 #homogenize baseline names

pl1=ggplot(resultat,aes(x=Year,y=value_moy,color=as.factor(func)))+geom_line(
)+facet_wrap(~growthlabel,ncol=4,scales="free")+
```

```
theme_bw()+theme(panel.grid=element_blank(),legend.position="none",
plot.title=element_text(size=14,face="bold"),strip.background=element_rect(fi
ll="white",color=NA))+ggtitle("a")+scale_color_brewer(palette =
"Set1")+ylab("Abundance")+scale_x_continuous(breaks=c(1,10,20,30,40))

pl1b=ggplot(resultat,aes(x=Year,y=value_moy,color=as.factor(func)))+geom_line
()+facet_wrap(~growthlabel,ncol=4,scales="free")+
theme_bw()+theme(panel.grid=element_blank(),legend.position="none",
plot.title=element_text(size=14,face="bold"),strip.background=element_rect(fi
ll="white",color=NA))+ggtitle("b")+scale_color_brewer(palette =
"Set1")+ylab("Abundance")+scale_y_log10(breaks =
scales::trans_breaks("log10", function(x) 10^x),labels =
scales::trans_format("log10", scales::math_format(10^.x)))+
annotation_logticks(sides = 'l')+scale_x_continuous(breaks=c(1,10,20,30,40))

pl2=ggplot(subset(resultat2,baseline==1),aes(x=as.factor(func),y=mean,color=a
s.factor(func)))+geom_boxplot()+facet_wrap(~growthlabel,scales="free",ncol=4)
+theme_bw()+
geom_hline(aes(yintercept=log(1+growth)),linetype="dashed",color="black")+the
me(panel.grid=element_blank(),plot.title=element_text(size=14,face="bold"),le
gend.position="none",
strip.background=element_rect(fill="white",color=NA))+
labs(color="Shape of population
dynamics")+ggtitle("c")+scale_color_brewer(palette = "Set1")+xlab("Shape of
population dynamics")+ylab("log(growth rate)")

pl3=ggplot(resultat2,aes(x=as.factor(baseline),y=mean,color=as.factor(func)))
+geom_boxplot()+
facet_wrap(~growthlabel,scales="free",ncol=4)+theme_bw()+
geom_hline(aes(yintercept=log(1+growth)),linetype="dashed",color="black")+the
me(panel.grid=element_blank(),plot.title=element_text(size=14,face="bold"),le
gend.position="bottom",
strip.background=element_rect(fill="white",color=NA))+
labs(color="Shape of population
dynamics")+ggtitle("d")+scale_color_brewer(palette = "Set1")+xlab("Baseline
year used to calculate the population trend")+ylab("log(growth rate)")

setwd(dir=filepath)
pdf("figure_4.pdf",width=7,height=11)
plot_grid(pl1,pl1b,pl2,pl3,ncol=1,rel_heights=c(1,1,1,1.4))
dev.off();
```

#### 5.3 Predict, Figure 5a-b

```
setwd(dir=filepath)
#build a prediction table:
setwd(dir="C:/Users/Duchenne/Documents/baseline/for_review/data_needed_to_run
_script")
tabtr=fread("tab_trend.txt")
tabtr$dat_site=paste(tabtr$Datasource,tabtr$Site)
pol=as.data.frame(poly(tabtr$baseline_trunc,3))
```

```
pol=round(pol,digits=11)
names(pol)[1:3]=paste0("p",names(pol)[1:3])
tabtr=cbind(tabtr,pol)
tabtr$Habitat_num=as.numeric(factor(tabtr$Habitat,levels=sort(unique(tabtr$Ha
bitat))))
tabtr$Continent_num=as.numeric(factor(tabtr$Continent,levels=sort(unique(tabt
r$Continent))))
tabtr$Study_num=as.numeric(factor(tabtr$Study,levels=sort(unique(tabtr$Study)
)))
tabtr$Order_num=as.numeric(factor(tabtr$Order,levels=sort(unique(tabtr$Order)
)))
tabtr$Datasource_num=as.numeric(factor(tabtr$Datasource,levels=sort(unique(ta
btr$Datasource))))
tabtr$dat_site_num=as.numeric(factor(tabtr$dat_site,levels=sort(unique(tabtr$
dat_site))))
tabtr$baseline_year=tabtr$baseline_trunc-min(tabtr$baseline_trunc)+1
tabtr$time_series_id_num=as.numeric(factor(tabtr$time_series_id,levels=sort(u
nique(tabtr$time_series_id))))
tabtr=subset(tabtr,nbann_pos>1 & nbann>=3)
variables=unique(tabtr[,c("baseline_trunc","p1","p2","p3","Continent_num","Ha
bitat_num","Continent","Habitat","baseline_year")])
rm(p1)
rm(p2)
rm(p3)
#function to predict:
predict_hglm=function(modelchains,nsampling,pred_table){
x=sample(1:nrow(obj),nsampling,replace=F)
attach(pred_table)
nbpred=nrow(pred_table)
fit=list()
colname=attributes(modelchains)$dimnames[[2]]
for (i in 1:nbpred){
fit[[i]] <-
c(#modelchains[x,paste0("intercept[",Continent_num[i],",",Habitat_num[i],"]")
]+apply(as.data.frame(modelchains[x,grep("Study",colname,fixed=T)]),1,mean)+

modelchains[x,paste0("baseline[",Continent_num[i],",",Habitat_num[i],",",base
line_year[i],"]")])
}

return(cbind(pred_table,data.frame(fit=sapply(fit,mean),media=sapply(fit,mean
),lwr=sapply(fit,quantile,prob=0.025),upr=sapply(fit,quantile,prob=0.975))))
}

#make prediction:
setwd(dir=filepath)
load("modelchains_withoutunique2.RData")
pred=predict_hglm(obj,1000,variables)
pred[,c("fit","lwr","upr")]=pred[,c("fit","lwr","upr")]
#plot them:
pl1b=ggplot(data=as.data.frame(pred),aes(x=baseline_trunc,y=fit,color=as.fact
```

```
or(Habitat),fill=as.factor(Habitat)))+
facet_wrap(~as.factor(Continent))+geom_errorbar(data=as.data.frame(pred),aes(
ymin=lwr,ymax=upr),alpha=0.2,width=0)+geom_point()+
theme_bw()+xlab("Baseline year")+ylab("Effect of the baseline year on
log(growth rate)\n relative to baseline year =
1970")+ggtitle("a")+labs(color="Habitat",fill="Habitat")+
scale_color_manual(values=c("dodgerblue3","tan4"))+scale_fill_manual(values=c
("dodgerblue3","tan4"))+theme(plot.title =
element_text(face="bold",size=16),axis.text=element_text(size=11),axis.title=
element_text(size=16),legend.text=element_text(size=16),legend.title=element_
text(size=16),strip.text= element_text(size=16),panel.grid =
element_blank(),strip.background = element_blank(),panel.border =
element_blank(), axis.line.x.bottom = element_line(color = 'black'),
    axis.line.y.left   = element_line(color =
'black'))+geom_hline(yintercept=0,linetype="dashed")+scale_y_continuous()

pl2b=ggplot(data=tabtr,aes(x=baseline_trunc,color=Habitat,fill=Habitat))+
geom_bar(adjust=2,position="stack")+
theme_bw()+xlab("Baseline year")+ylab("Number of abundance
trends")+ggtitle("b")+labs(color="Habitat",fill="Habitat")+
scale_color_manual(values=c("dodgerblue3","tan4"))+scale_fill_manual(values=c
("dodgerblue3","tan4"))+
theme(plot.title =
element_text(face="bold",size=14),axis.text=element_text(size=11),
axis.title=element_text(size=16),legend.text=element_text(size=16),legend.tit
le=element_text(size=16),strip.text=
element_text(size=16),panel.grid=element_blank(),strip.background =
element_blank(),panel.border = element_blank(), axis.line.x.bottom =
element_line(color = 'black'),
axis.line.y.left   = element_line(color = 'black'))+
facet_wrap(~Continent)+scale_x_continuous(breaks=seq(1970,2000,10))

setwd(dir=filepath)
pdf("figure_5.pdf",width=10,height=10)
plot_grid(pl1b,pl2b,ncol=1,align="v")
dev.off();
```

#### 5.4 Variance decomposition, Figure 6a

```
setwd(dir=filepath)
#Fig 6a:
suma=fread("summary_model_withoutunique2.txt")
b=suma[grep("sd.",suma$varia,fixed=T),]
#translate variable names:
b$varia2=NA
b$varia2[b$varia=="sd.s"]="Site"
b$varia2[b$varia=="sd.o"]="Order"
b$varia2[b$varia=="sd.a"]="Data source"
b$varia2[b$varia=="sd.ts"]="Time series ID"
b$varia2[b$varia=="sd.resi"]="Residual"
b$varia2[b$varia=="sd.fixed"]="Fixed Effects"
b$mean[b$varia=="sd.baseline"]=apply(suma[grep("baseline[",suma$varia,fixed=T
```

```
),"mean"],2,sd)
b$varia2[b$varia=="sd.baseline"]="Baseline"
#calculate variance ratio from standard deviations:
tabtr=fread("tab_trend.txt")
b$variance=(b$mean)/(sum(b$mean))

b=b[order(b$variance,decreasing=T),]
b$varia2=factor(b$varia2,levels=b$varia2)
pl3b=ggplot(data=b,
aes(x=varia2,y=variance))+geom_bar(stat="identity",fill="black")+
ylab("Percentage of explained variation\nin abundance trends")+
theme_bw()+
theme(plot.title =
element_text(face="bold",size=14),axis.text=element_text(size=11),axis.text.x
=element_text(size=11,angle=45,hjust=1),axis.title=element_text(size=14),axis
.title.x=element_blank(),legend.text=element_text(size=14),legend.title=eleme
nt_text(size=14),strip.text=
element_text(size=14),panel.grid=element_blank(),legend.position="none")+ggti
tle("b")+scale_y_continuous(labels = scales::percent)
```

#### 5.5 Figure 6b&c

```
setwd(dir=filepath)
suma=fread("summary_model_withoutunique2.txt")
tabtr$Study_num=as.numeric(factor(tabtr$Study,levels=sort(unique(tabtr$Study)
)))
bidon=unique(tabtr[,c("Study","Study_num")])
bidon=bidon[order(bidon$Study_num),]
studies=cbind(bidon,suma[grep("Study",suma$varia,fixed=F),])
studies$Study[1]=paste0(studies$Study[1]," (ref.)")
studies=studies[,c("Study","mean","l95","u95")]
studies$type="model estimate"

#EMPIRICAL AVERAGE BEFORE CORRECTION BY MODEL
tabtr=fread("tab_trend.txt")
#take only the originale time series
tabtr= tabtr %>% group_by(time_series_id) %>%
mutate(base=min(baseline_trunc))
tabtr3=subset(tabtr,baseline_trunc==base & nbann_pos>1)
#centered on the reference study (BIOTIME)
tabtr3$trend=tabtr3$trend-mean(tabtr3$trend[tabtr3$Study=="BIOTIME"])
b=Rmisc::group.CI(trend~Study,data=tabtr3)
b=b[,c(1,3,2,4)]
b$Study[b$Study=="BIOTIME"]="BIOTIME (ref.)"
b$type="empirical mean"
names(b)=names(studies)

studies=rbind(studies,b)

pl1c=ggplot(data=subset(studies,type=="model estimate"),
aes(x=Study,y=mean))+geom_point()+
```

```
geom_errorbar(aes(ymin=l95,ymax=u95),width=0)+
geom_hline(yintercept=0,linetype="dashed")+
ylab("Dataset effect on estimated abundance trend\n(relatively to the
reference level)")+
theme_bw()+
theme(plot.title =
element_text(face="bold",size=14),axis.text=element_text(size=11),axis.title=
element_text(size=14),axis.title.y=element_blank(),legend.text=element_text(s
ize=14),legend.title=element_text(size=14),strip.text=
element_text(size=14),panel.grid=element_blank(),legend.position="none")+coor
d_flip()+
ggtitle("c")

####VARIANCE AND DURATION:
tabtr=fread("tab_trend.txt")
tabtr$dat_site=paste(tabtr$Datasource,tabtr$Site)
pol=as.data.frame(poly(tabtr$baseline_trunc,3))
pol=round(pol,digits=11)
names(pol)[1:3]=paste0("p",names(pol)[1:3])
tabtr=cbind(tabtr,pol)
tabtr$Habitat_num=as.numeric(factor(tabtr$Habitat,levels=sort(unique(tabtr$Ha
bitat))))
tabtr$Continent_num=as.numeric(factor(tabtr$Continent,levels=sort(unique(tabt
r$Continent))))
tabtr$Study_num=as.numeric(factor(tabtr$Study,levels=sort(unique(tabtr$Study)
)))
tabtr$Order_num=as.numeric(factor(tabtr$Order,levels=sort(unique(tabtr$Order)
)))
tabtr$Datasource_num=as.numeric(factor(tabtr$Datasource,levels=sort(unique(ta
btr$Datasource))))
tabtr$dat_site_num=as.numeric(factor(tabtr$dat_site,levels=sort(unique(tabtr$
dat_site))))
tabtr$time_series_id_num=as.numeric(factor(tabtr$time_series_id,levels=sort(u
nique(tabtr$time_series_id))))
tabtr$baseline_year=tabtr$baseline_trunc-min(tabtr$baseline_trunc)+1
tabtr$trend_error=tabtr$sd
tabtr=subset(tabtr,nbann_pos>1 & nbann>=3)
variables=unique(tabtr[,c("Study_num","nbann","Study","trend_error")])
load("modelchains_withoutunique2.RData")
#function to predict variance:
rm(modelchains)
rm(pred_table)
rm(nsampling)
predict_hglm_var=function(modelchains,nsampling,pred_table){
x=sample(1:nrow(obj),nsampling,replace=F)
attach(pred_table)
nbpred=nrow(pred_table)
fit=list()
```

```
colname=attributes(modelchains)$dimnames[[2]]
for (i in 1:nbpred){
fit[[i]] <-
exp(modelchains[x,paste0("intercept_dispersion[",Study_num[i],"]")]+modelchai
ns[x,"error_effect"]*trend_error[i]+modelchains[x,paste0("nbann[",Study_num[i
],"]")]*nbann[i]+modelchains[x,paste0("nbann2[",Study_num[i],"]")]*(nbann[i]^
2))
}

return(cbind(pred_table,data.frame(fit=sapply(fit,mean),media=sapply(fit,mean
),lwr=sapply(fit,quantile,prob=0.025),upr=sapply(fit,quantile,prob=0.975))))
}

#predict variance
pred=predict_hglm_var(obj,1000,variables)

#function to extract residuals:
residuals_hglm=function(modelchains,nsampling,pred_table){
x=sample(1:nrow(modelchains),nsampling,replace=F)
attach(pred_table)
nbpred=nrow(pred_table)
fit=c()
colname=attributes(modelchains)$dimnames[[2]]
for (i in 1:nbpred){
fit[i] <-pred_table$trend[i]-
mean(c(modelchains[x,paste0("intercept[",Continent_num[i],",",Habitat_num[i],
"]")]+modelchains[x,paste0("Study[",Study_num[i],"]")]+modelchains[x,paste0("
baseline[",Continent_num[i],",",Habitat_num[i],",",baseline_year[i],"]")]+
modelchains[x,paste0("Order[",Order_num[i],"]")]+modelchains[x,paste0("Dataso
urce[",Datasource_num[i],"]")]+
modelchains[x,paste0("dat_site[",dat_site_num[i],"]")]+modelchains[x,paste0("
time_series_id[",time_series_id_num[i],"]")]))
}
return(cbind(pred_table,data.frame(residuals=fit)))
}
set.seed(3)

resid=residuals_hglm(obj,1000,tabtr)
#observed variance in residuals
bidon=resid %>% group_by(nbann,Study) %>% summarise(variance=var(residuals))
#plot them:
pl2c=ggplot(data=as.data.frame(pred),aes(x=nbann,y=fit,col=Study))+
geom_line()+
theme_bw()+xlab("Number of year with data in the time
series")+ylab("\nResidual variance in abundance
trends")+labs(color="Dataset")+
theme(plot.title =
element_text(face="bold",size=14),axis.text=element_text(size=11),axis.title=
element_text(size=14),legend.text=element_text(size=14),legend.title=element_
text(size=14),strip.text= element_text(size=14),panel.grid =
element_blank())+geom_point(data=bidon,aes(x=nbann,y=variance),pch=21,fill=NA
```

```
)+
scale_y_continuous(trans = "log",breaks=c(0.001,0.0001,0.01,0.1,1,10),labels
= trans_format("log10",math_format(10^.x)))
pl2cb=ggplot(data=tabtr,aes(x=nbann,fill=Study,col=Study))+geom_bar(position=
"stack")+theme_void()+
theme(plot.title =
element_text(face="bold",size=14),axis.text.y=element_text(size=8),axis.title
=element_blank(),axis.title.y=element_blank(),legend.text=element_blank(),leg
end.title=element_blank(),strip.text=
element_blank(),panel.grid=element_blank(),legend.position="none",axis.line.y
=
element_line(),axis.ticks.y=element_line(),axis.ticks.length=unit(0.1,"cm"))+
ggtitle("a")+scale_y_continuous(n.breaks=3)

pp=plot_grid(pl2cb,pl2c,ncol=1,align="v",axis="lr",rel_heights=c(0.3,1))

pp2=plot_grid(pl3b,pl1c,ncol=1,align="v",axis="lr",rel_heights=c(1,1))

pdf("figure_6.pdf",width=12,height=8)
gridExtra::grid.arrange(pp,pp2,ncol=2,widths=c(1.2,1))
dev.off();
```

### 6 Supplementary

#### 6.1 Figure S1 & Table S1

```
setwd(dir=filepath)
tabf=fread("compiled_dataset.txt")
tabf$Rank[tabf$Rank=="CLASS"]="ORDER"
tabf$Study[tabf$Study=="Klink"]="van Klink"
bidon=tabf %>% dplyr::group_by(Study,Order) %>%
dplyr::summarise(value=n_distinct(time_series_id))

pl1=ggplot(data=bidon,aes(y=Order,x=value,fill=Study,col=Study))+theme_bw()+
geom_bar(stat="identity",position =
position_dodge2(preserve="single"))+ggtitle("a")+ylab("Taxonomic groups")+
theme(axis.text.x = element_text(angle = 0, hjust =
0),legend.position="bottom")+xlab("Number of time series")+
scale_x_log10(breaks = scales::trans_breaks("log10", function(x) 10^x),labels
= scales::trans_format("log10",
scales::math_format(10^.x)))+annotation_logticks(sides = "l", outside =
TRUE)+labs(fill="",color="")

bidon=tabf %>% dplyr::group_by(Study,Rank) %>%
dplyr::summarise(nb=n_distinct(time_series_id))

pl2=ggplot(data=bidon,aes(x=Study,y=nb,fill=Rank))+theme_bw()+
geom_bar(stat="identity")+ggtitle("b")+xlab("Dataset")+
theme(axis.text.x = element_text(angle = 45, hjust =
```

```
1),legend.text=element_text(size=6),legend.position="bottom")+ylab("Number of
time
series")+guides(fill=guide_legend(nrow=2,byrow=TRUE),color=guide_legend(nrow=
2,byrow=TRUE))+
scale_fill_brewer(palette="Set3")+labs(fill="")

leg1 <- get_legend(pl1)
leg1=as_ggplot(leg1)
pl1=pl1+theme(legend.position="none")
leg2 <- get_legend(pl2)
leg2=as_ggplot(leg2)
pl2=pl2+theme(legend.position="none")

setwd(dir=filepath)
png("figure_S1.png",width=1200,height=1700,res=180)
grid.arrange(pl1,pl2,leg1,leg2,ncol=2,widths=c(2,1),heights=c(4,0.5))
dev.off();

bidon=tabf %>% dplyr::group_by(Study,Continent,Habitat) %>%
dplyr::summarise(value=n_distinct(time_series_id))
bidon=dcast(bidon,Study+Continent~Habitat)
fwrite(bidon,"TableS1.txt")
```

#### 6.2 Figure S4

```
set.seed(22)
bidon=data.frame(x=1:50,y=c(rnorm(50,10,1)-seq(0,0.1*50,length.out
=50),rnorm(50,10,0.1)-seq(0,0.01*50,length.out
=50)),sp=rep(c("sp1","sp2"),each=50))

pl1=ggplot(data=bidon,aes(x=x,y=y,col=sp,fill=sp))+theme_bw()+geom_point()+st
at_smooth(method="lm",alpha=0.1)+xlab("Time (Years)")+ylab("Abundance")+
labs(color="Species",fill="Species")+ggtitle("a")+scale_color_manual(values=c
("darkolivegreen","darkmagenta"))+scale_fill_manual(values=c("darkolivegreen"
,"darkmagenta"))+theme(panel.grid=element_blank())

bidon=bidon %>% group_by(sp) %>% mutate(y2=scale(y))
pl2=ggplot(data=bidon,aes(x=x,y=y2,col=sp,fill=sp))+theme_bw()+geom_point()+s
tat_smooth(method="lm",alpha=0.1)+xlab("Time (Years)")+ylab("Abundance
standardized at the time series level")+
labs(color="Species",fill="Species")+ggtitle("b")+scale_color_manual(values=c
("darkolivegreen","darkmagenta"))+scale_fill_manual(values=c("darkolivegreen"
,"darkmagenta"))+theme(panel.grid=element_blank())

setwd(dir=filepath)
png("figure_S4.png",width=1000,height=2000,res=170)
grid.arrange(pl1,pl2,ncol=1)
dev.off();
```

#### 6.3 Figure S5

```
setwd(dir=filepath)
tabtr=fread("tab_trend.txt")
tabtr$cat="1"
tabtr$cat[tabtr$nbann_pos>1]=">1"
tabtr=subset(tabtr,nbann>=3)

pl1=ggplot(data=tabtr,aes(x=abs(trend),color=cat,fill=cat))+geom_density(alph
a=0.4)+
scale_color_manual(values=c("firebrick4","chartreuse4"))+scale_fill_manual(va
lues=c("firebrick4","chartreuse4"))+theme_bw()+
theme(plot.title =
element_text(face="bold",size=14),axis.text=element_text(size=11),axis.title=
element_text(size=14),legend.text=element_text(size=14),legend.title=element_
text(size=14),strip.text= element_text(size=14),panel.grid =
element_blank())+guides(color = guide_legend(override.aes = list(size = 2) )
)+xlab(expression( "|log(growth rate)|"))+ylab("Density")+
scale_x_continuous(trans="sqrt",breaks=c(0,1,10,100,200,300))+labs(fill="n of
positive\nabundance estimates",color="n of positive\nabundance estimates")

png("figure_S5.png",width=1200,height=800,res=170)
pl1
dev.off();
```

#### 6.4 Figure S7

```
setwd(dir=filepath)
load("simues.RData")
resultat=res[[1]]
resultat2=res[[2]]
resultat$growthlabel=factor(paste0("r = ",1+resultat$growth),c("r = 1","r =
0.95","r = 0.9","r = 0.85"))
resultat2$growthlabel=factor(paste0("r = ",1+resultat2$growth),c("r = 1","r =
0.95","r = 0.9","r = 0.85"))
resultat2$baseline[resultat2$baseline==0]=1 #homogenize baseline names

# compute lower and upper whiskers
ylimdf=subset(resultat2,growth==(-0.15)) %>% group_by(baseline,func) %>%
summarise(limi1=quantile(mean, prob=0.00025),limi2=quantile(mean,
prob=0.975))
ylim1 =c(min(ylimdf$limi1),max(ylimdf$limi2))

ps7=ggplot(subset(resultat2,growth==(-
0.15)),aes(x=as.factor(baseline),y=mean,color=as.factor(func)))+geom_boxplot(
outlier.shape = NA)+
facet_wrap(~growthlabel,scales="free",ncol=1)+theme_bw()+
geom_hline(aes(yintercept=log(1+growth)),linetype="dashed",color="black")+the
me(panel.grid=element_blank(),plot.title=element_text(size=14,face="bold"),le
gend.position="bottom",
strip.background=element_rect(fill="white",color=NA))+
```

```
labs(color="Shape of population dynamics")+scale_color_brewer(palette =
"Set1")+xlab("Baseline year used to calculate the population
trend")+ylab("log(growth rate)")+coord_cartesian(ylim = ylim1*2)

setwd(dir=filepath)
png("figure_S7.png",width=1000,height=1300,res=150)
ps7
dev.off();
```

##### 6.5 Figure S8

```
setwd(dir=filepath)
load("modelchains_withoutunique2.RData")
colname=attributes(obj)$dimnames[[2]]
suma=as.data.frame(obj[(nrow(obj)-
2000):nrow(obj),grep("Order",colname,fixed=T)])
suma=melt(suma)
suma$Order_num=gsub("]","",substr(suma$variable,7,8),fixed=T)

tabtr=fread("tab_trend.txt")
tabtr=subset(tabtr,nbann_pos>1 & nbann>=3)
tabtr$Order_num=as.numeric(factor(tabtr$Order,levels=sort(unique(tabtr$Order)
)))
bidon=unique(tabtr[,c("Order","Order_num")])

suma=merge(suma,bidon,by="Order_num")
bidon=suma %>% dplyr::group_by(Order) %>% summarise(moy=mean(value))
bidon=bidon[order(bidon$moy),]
suma$Order=factor(suma$Order,bidon$Order)

pl1=ggplot(as.data.frame(suma), aes(x=value, y=Order))+
geom_vline(xintercept=0,linetype="dashed")+
stat_summary(fun="mean",fun.min = function(z) { quantile(z,0.025) },fun.max =
function(z) { quantile(z,0.975) })+
theme_bw()+
ggtitle("")+
theme(plot.title =
element_text(face="bold",size=14),axis.text=element_text(size=11),axis.title=
element_blank(),legend.text=element_text(size=14),legend.title=element_text(s
ize=14),strip.text=
element_text(size=14),strip.background=element_blank(),legend.position="none"
,panel.grid=element_blank())

setwd(dir=filepath)
png("figure_S8.png",width=1000,height=1300,res=150)
pl1
dev.off();
```

#### 6.6 Figure S9

```
setwd(dir=filepath)
liste=fread("linearity_time_series.txt")
tabtr=fread("tab_trend.txt")
tabtr=subset(tabtr,nbann_pos>1 & nbann>=3)
tabtr=merge(tabtr,liste[,c("time_series_id","abond_moy")],by="time_series_id"
)

pl1=ggplot(data=tabtr,aes(x=abond_moy,y=trend,color=Habitat))+geom_point(size
=0.9,alpha=0.4)+facet_wrap(~Study,scales="free")+
scale_color_manual(values=c("dodgerblue3","tan4"))+scale_fill_manual(values=c
("dodgerblue3","tan4"))+theme_bw()+theme(plot.title =
element_text(face="bold",size=14),axis.text=element_text(size=11),axis.title=
element_text(size=14),legend.text=element_text(size=14),legend.title=element_
text(size=14),strip.text= element_text(size=14),panel.grid =
element_blank())+guides(color = guide_legend(override.aes = list(size = 2) )
)+ylab("Abundance trends")+xlab("Average abundance of the original time
series")+
scale_x_log10(breaks = scales::trans_breaks("log10", function(x) 10^x),labels
= scales::trans_format("log10", scales::math_format(10^.x)))+
annotation_logticks(sides = 'b')

setwd(dir=filepath)
png("figure_S9.png",width=1300,height=1000,res=150)
pl1
dev.off();
```
